# Inhibition of HSP90 distinctively modulates the global phosphoproteome of *Leishmania mexicana* developmental stages

**DOI:** 10.1101/2023.07.19.549707

**Authors:** Exequiel O. J. Porta, Liqian Gao, Paul W. Denny, Patrick G. Steel, Karunakaran Kalesh

**Affiliations:** Department of Chemistry, Durham University, Durham, United Kingdom; School of Pharmaceutical Sciences, Shenzhen Campus of Sun Yat-sen University, Shenzhen, China; Department of Biosciences, Durham University, Durham, United Kingdom; School of Health and Life Sciences, Teesside University, Middlesbrough, United Kingdom; National Horizons Centre, Darlington, United Kingdom

**Keywords:** Phosphorylation, *Leishmania*, Protein kinases, HSP90, TMT labelling, LC-MS/MS, RNA helicase

## Abstract

Heat shock protein 90 (HSP90) is an evolutionary conserved chaperone protein that plays a central role in the folding and maturation of a large array of client proteins. In the unicellular parasite *Leishmania*, the etiological agent of the neglected tropical disease leishmaniasis, treatment of the classical HSP90 inhibitor tanespimycin leads to dose- and time-dependent differentiation from promastigote to amastigote stage, eventually culminating in parasite killing. Although this suggests a crucial role of the HSP90 in the life cycle control of *Leishmania*, the underlying molecular mechanism remains unknown. Using a combination of phosphoproteome enrichment and tandem mass tag (TMT) labelling-based quantitative proteomic mass spectrometry (MS), we robustly identified and quantified 1,833 phosphorylated proteins across three life cycle stages of *Leishmania mexicana* (*L. mexicana*) parasite. Protein kinase domain was the most enriched protein domain in the *L. mexicana* phosphoproteome. Additionally, this study systematically characterised the perturbing effect of HSP90 inhibition on the global phosphoproteome of *L. mexicana* across its life cycle stages and showed that the tanespimycin treatment causes substantially distinct molecular effects in promastigote and amastigote forms. Whilst phosphorylation of HSP90 and its co-chaperon HSP70 was decreased in amastigotes, the opposite effect was observed in promastigotes. Additionally, our results showed that while kinase activity and microtubule motor activity are highly represented in the negatively affected phosphoproteins of the promastigotes, whereas ribosomal proteins, protein folding, and proton channel activity are preferentially enriched in the perturbed amastigote phosphoproteome. Our results also show that RNA helicase domain was distinctively enriched among the positively affected RNA-binding amastigote phosphoproteome. This study reveals the dramatically different ways the HSP90 inhibition stress modulates the phosphoproteome of the pathogenic amastigotes and provides in-depth insight into the scope of selective molecular targeting in the therapeutically relevant amastigote forms.

**IMPORTANCE:** In the unicellular parasites *Leishmania* spp., the etiological agents of leishmaniasis, a complex infectious disease that affects 98 countries in 5 continents, chemical inhibition of HSP90 protein, a master regulator of protein homeostasis, leads to differentiation from promastigote to amastigote stage, eventually culminating in parasite death. However, the underlying molecular mechanism remains unknown. Recent studies suggest a fundamentally important role of RNA-binding proteins (RBPs) in regulating the downstream effects of the HSP90 inhibition in *Leishmania*. Phosphorylation-dephosphorylation dynamics of RNA-binding proteins (RBPs) in higher eukaryotes serves as an important on/off switch to regulate RNA processing and decay in response to extracellular signals and cell cycle check points. In the current study, using a combination of highly sensitive tandem mass tag (TMT) labelling-based quantitative proteomic mass spectrometry (MS) and robust phosphoproteome enrichment, we show for the first time that the HSP90 inhibition distinctively modulates global protein phosphorylation landscapes in the different life cycle stages of *Leishmania*, shedding light into a crucial role of the posttranslational modification in the differentiation of the parasite under HSP90 inhibition stress. This work provides insights into the importance of HSP90-mediated protein cross-talks and regulation of phosphorylation in *Leishmania*, thus significantly expanding our knowledge of the posttranslational modification in *Leishmania* biology.

---

*Leishmania* spp. are unicellular eukaryotic parasites that cause a complex spectrum of diseases in humans and animals (1). The parasite has a digenetic life cycle alternating between sandfly vectors and mammalian hosts adapting to changing environments via life cycle specific expression of genes. The regulation of gene expression in *Leishmania* spp. is mostly posttranscriptional and involves processes such as mRNA processing, mRNA decay, protein translation and posttranslational modifications (PTMs) (2). Amongst the different PTMs profiled during *Leishmania* spp. differentiation protein phosphorylation has been found to occur on proteins that correlate well to parasite differentiation through its life cycle, such as ribosomal proteins (RPs), cytoskeletal proteins (CSPs), heat shock proteins (HSPs), RNA-binding proteins (RBPs), protein kinases (PKs) and protein phosphatases (PPs) (3, 4). As the majority of *Leishmania* spp. phosphoproteins identified to date are differentially expressed in the different life cycle stages (3, 5–7), a complex network of protein phosphorylation and dephosphorylation events catalysed by life cycle-specific protein kinases and protein phosphatases is thought to play a crucial role in the elusive processes dictating *Leishmania* spp. differentiation.

HSP90 has been previously identified as a downstream client of phosphorylation-mediated signalling in *Leishmania* spp. (8). Interestingly, treatment of HSP90 inhibitors leads to dose- and time-dependent differentiation of *Leishmania* spp. promastigotes to amastigotes in axenic cultures, suggesting a central role for the HSP90 in the life cycle control of the organism (9–11). We have recently shown that inhibition of HSP90 using the classical inhibitor tanespimycin causes a repression of ribosomal protein synthesis in *L. mexicana* promastigotes (12). More recently, we have also shown that the inhibition of HSP90 causes widespread perturbation of RNA-protein interactions in both promastigote and amastigote life cycle stages of *L. mexicana* (13). The RNA interactions of a substantial portion of the *L. mexicana* protein kinome were perturbed by the HSP90 inhibition (13). Phosphorylation and dephosphorylation dynamics of RBPs by the coordinated action of PKs and PPs is an important regulatory mechanism to control RNA processing and decay in response to cell cycle checkpoints and extracellular signals (14). Therefore, we envisaged a large-scale study capturing the global phosphoproteome of *L. mexicana* and characterising its perturbations under the influence of the HSP90 inhibition. In order to accurately capture the modulations in the phosphoproteome, we combined for the first time the HSP90 inhibitor treatment in log-phase promastigote (LPP), stationary-phase promastigote (SPP) and axenic amastigote (AXA) life cycle stages of *L. mexicana* with phosphoproteome enrichment, followed by tandem mass tag (TMT) labelling (15) based quantitative proteomic mass spectrometry (MS). This study significantly expands the protein phosphorylation landscape of *Leishmania* spp., providing robust identification of several thousand of phosphorylation sites and the modulatory effect of the HSP90 inhibition across the three different life cycle stages of this protozoan parasite.

## RESULTS

### Global changes in protein phosphorylation of the three life cycle stages of *Leishmania mexicana*

TMT labelling-based quantitative proteomic MS of enriched phosphoproteome from LPP, SPP and AXA life cycle stages of *L. mexicana* was performed in three biological replicates. A total of 1,833 phosphoproteins were identified with a minimum of two unique peptides across the three life cycle stages (Table S1). As shown in the volcano plots, Fig. 1A and Fig. 1B, we observed quantitative differences in the global phosphoproteome patterns across the life cycle stages (Tables S2, S3 and S4). An increased phosphorylation of HSP90 and HSP70 in the amastigote stage of *Leishmania* compared to its promastigotes was recently reported (8). In agreement with these findings, our results show a quantitative increase in the phosphorylation of both proteins upon differentiation to AXA (Fig. 1A and Fig. 1B). A representative tandem mass spectrum of a phosphorylation site identification in the *L. mexicana* HSP90 is given in Fig. 1C. In addition to the previously reported phosphorylation at residues Thr_211_, Thr_216_, Ser_289_, Ser_371_, Ser_526_, Ser_594_ and Ser_595_ (8), our study identified five previously unknown phosphorylation sites at Ser_38_, Ser_48_, Thr_239_, Thr_256_ and Ser_477_ in the *L. mexicana* HSP90 (Supplementary Fig. S1). Furthermore, this large-scale study revealed the phosphorylation sites and quantitative changes in the global phosphoproteome including important signalling kinases, RBPs, motor proteins, hydrolases and ligases (Supplementary Fig. S2) across the three life cycle stages of *L. mexicana* (Tables S1 to S4).

**FIG. 1.**
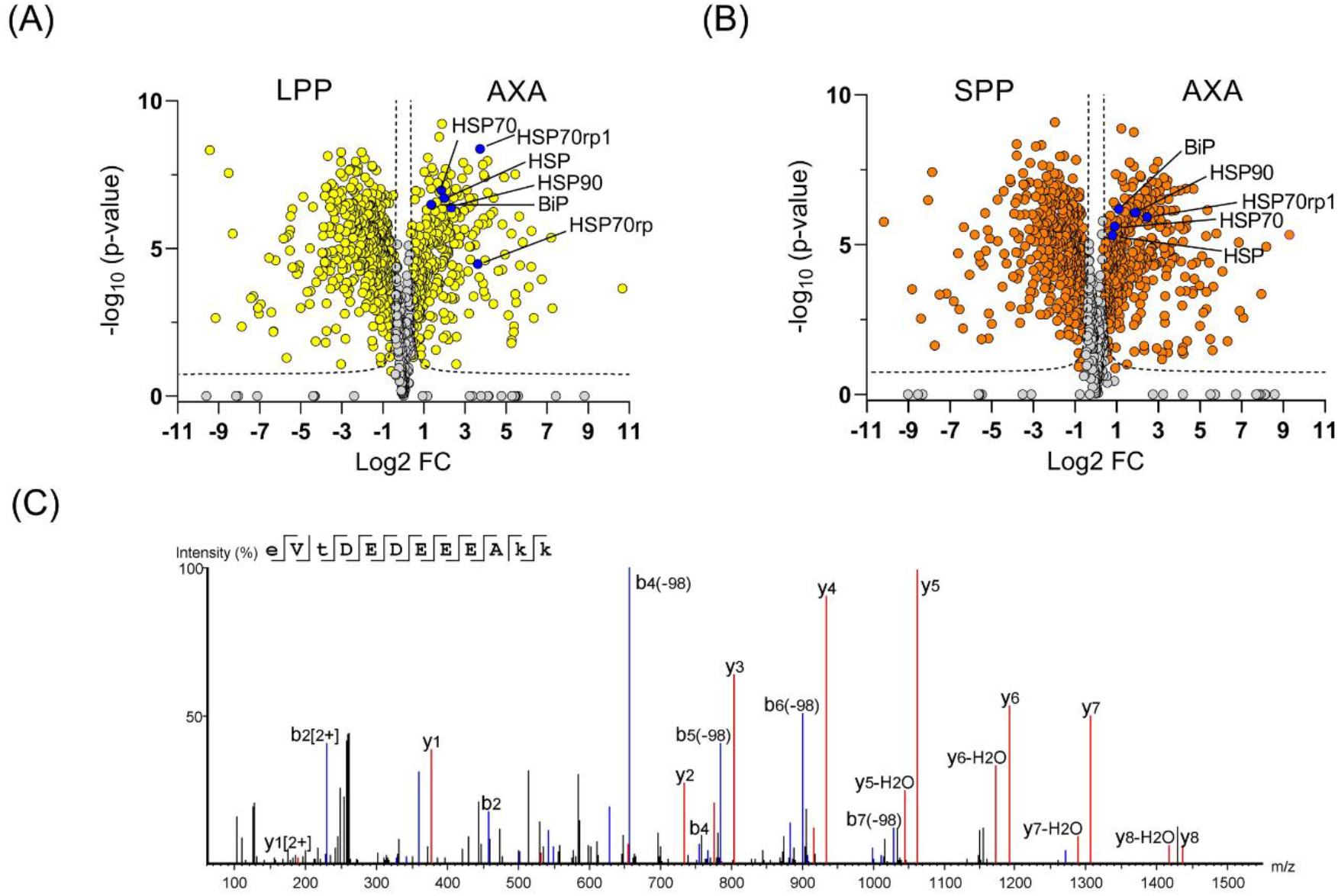
Global changes in the phosphoproteome of *L. mexicana* across its log-phase promastigote (LPP), stationary-phase promastigote (SPP) and axenic amastigote (AXA) life cycle stages profiled by phosphoproteome enrichment followed by tandem mass tag (TMT) labelling-based quantitative proteomic mass spectrometry (MS). All phosphoproteome enrichment experiments were performed in three biological replicates. (A) and (B) Volcano plots showing differential enrichment of phosphoproteins between LPP and AXA and SPP and AXA respectively. A modified *t* test with permutation-based FDR statistics (250 permutations, FDR = 0.05) was applied to compare the quantitative differences in the phosphoproteins between the life cycle groups. Heat shock proteins (HSPs) that showed increased phosphorylation in the AXA are highlighted in blue filled circles. (C) MS/MS spectrum of Thr_216_ phosphorylated peptide from the *L. mexicana* HSP90.

### Functional analysis of the *L. mexicana* phosphoproteins

Analysis of protein families and domains in the *L. mexicana* global phosphoproteome revealed protein kinase (PK) as the most enriched protein domain (Supplementary Fig. S3). PK phosphorylation was found to be differentially regulated across the *L. mexicana* life cycle stages (Supplementary Fig. S4). Classification of *L. mexicana* PKs according to their catalytic domain conservation (16) revealed the majority of PKs in the CMGC and STE groups phosphorylated (Fig. 2).

**FIG. 2.**
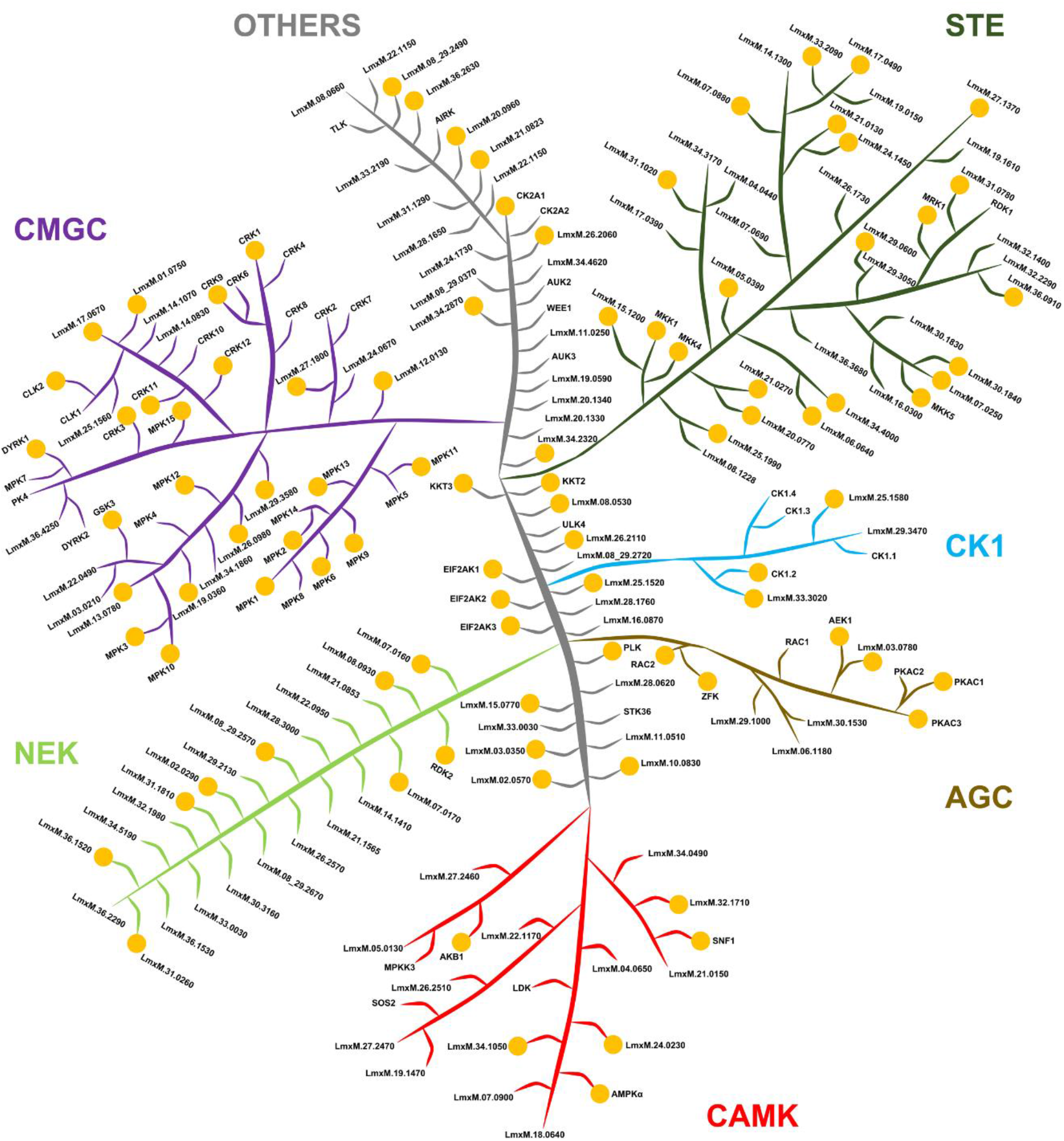
Phosphorylation in *L. mexicana* protein kinome. Classification of the protein kinases in *L. mexicana* according to their catalytic domain types namely GMGC, STE, NEK, CK1, AGC, CAMK and OTHERS. The image has no phylogenetic significance and is for illustrative purpose only. The orange dots represent protein kinases in which phosphorylation was detected in this study.

Next, in order to gain more insight into the functional differences between the enriched phosphoproteins of the promastigotes and the amastigotes, we performed Gene Ontology (GO) analyses of the statistically significant differentially expressed phosphoproteins. The most enriched Molecular Function (MF) GO terms of the three life cycle stages are shown in Fig. 3. In contrast to the two promastigote stages, which showed a preferential enrichment of PK activity, the amastigote stage showed binding interactions, particularly RNA binding and unfolded protein binding preferentially enriched amongst their phosphoproteins. Intriguing differences were also observed in the Biological Process (BP) GO terms of the phosphoproteins of the three life cycle stages (Supplementary Fig. S5). In the LPP, the most enriched BP term was protein phosphorylation. However, in the SSP along with protein phosphorylation, cytoplasmic translation was found to be a prominent BP. In a continuum, in the AXA, the translation became the most enriched BP term of the phosphoproteins. Similarly, striking differences were also observed in the Cellular Component (CC) GO terms of the phosphoproteins between the promastigotes and the amastigotes (Supplementary Fig. S6). In both LPP and SSP, cilium and axoneme were the most enriched CC GO terms, suggesting potential involvement of protein phosphorylation in the structure and function of flagellum and plausible reliance of the parasite cell motility on phosphorylation in the promastigotes. In contrast, enrichment of the supramolecular complex CC term in the AXA phosphoproteins suggests a life cycle-selective role for protein phosphorylation in the formation and function of large, multi-protein complexes involved in cellular processes such as protein translation.

**FIG. 3.**
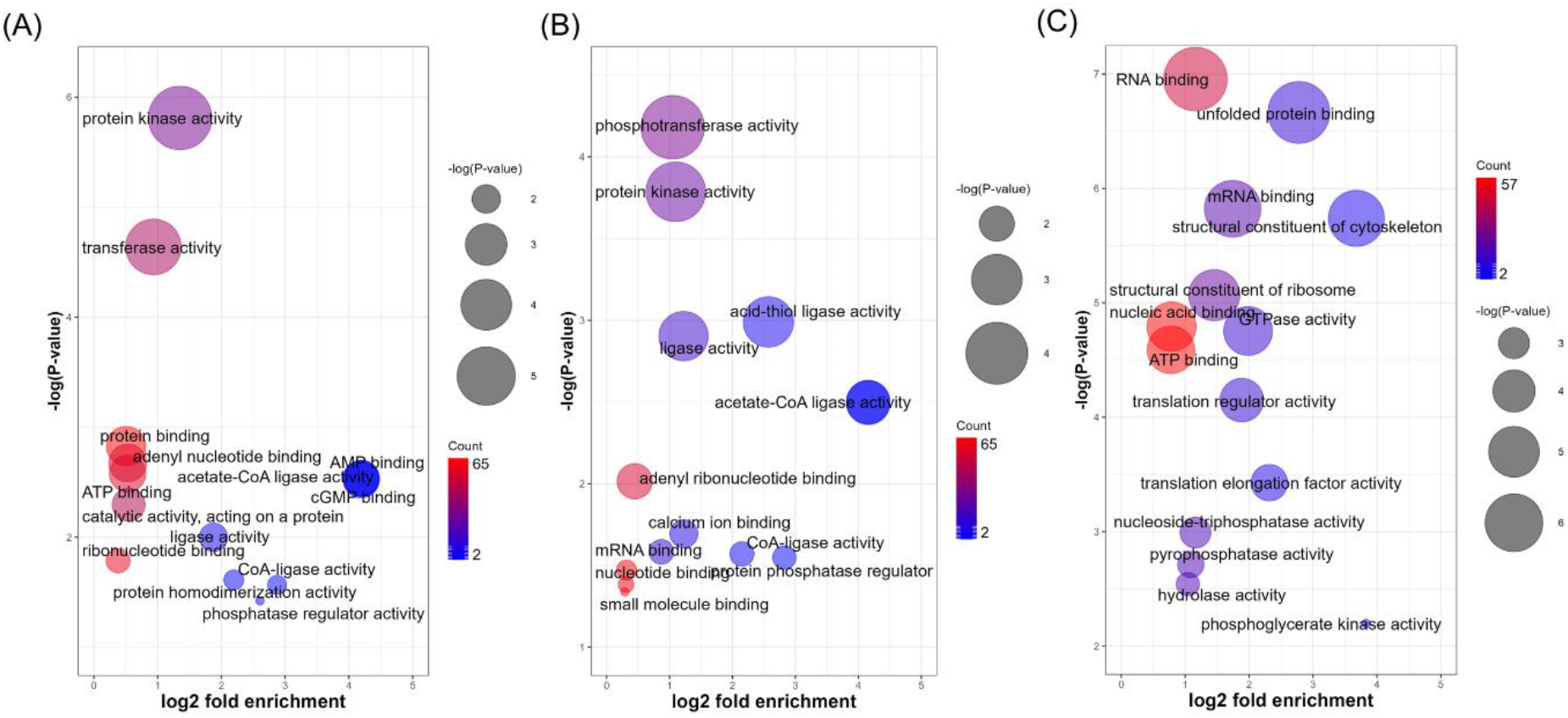
Molecular Function (MF) gene ontology (GO) terms enriched in (A) LPP, (B) SPP and (C) AXA phosphoproteins.

### Physicochemical properties of the *L. mexicana* global phosphoproteome

We then compared the physicochemical properties hydrophobicity, isoelectric point and molecular weight of the *L. mexicana* phosphoproteome (Fig. 4A, Fig. 4B, Supplementary Fig. S7). Cumulative distributions of the physicochemical properties in the LPP, SPP and AXA phosphoproteins and the entire *L. mexicana* proteome revealed that the proteins undergoing phosphorylation in all life cycle stages are generally hydrophilic (Fig. 4C, Supplementary Fig. S7C). The analysis also revealed that the LPP and SPP phosphorylation substrates are larger proteins (Fig. 4D, Supplementary Fig. S7D), and the AXA phosphorylation substrates have comparatively more acidic isoelectric points (Fig. 4E, Supplementary Fig. S7E).

**FIG. 4.**
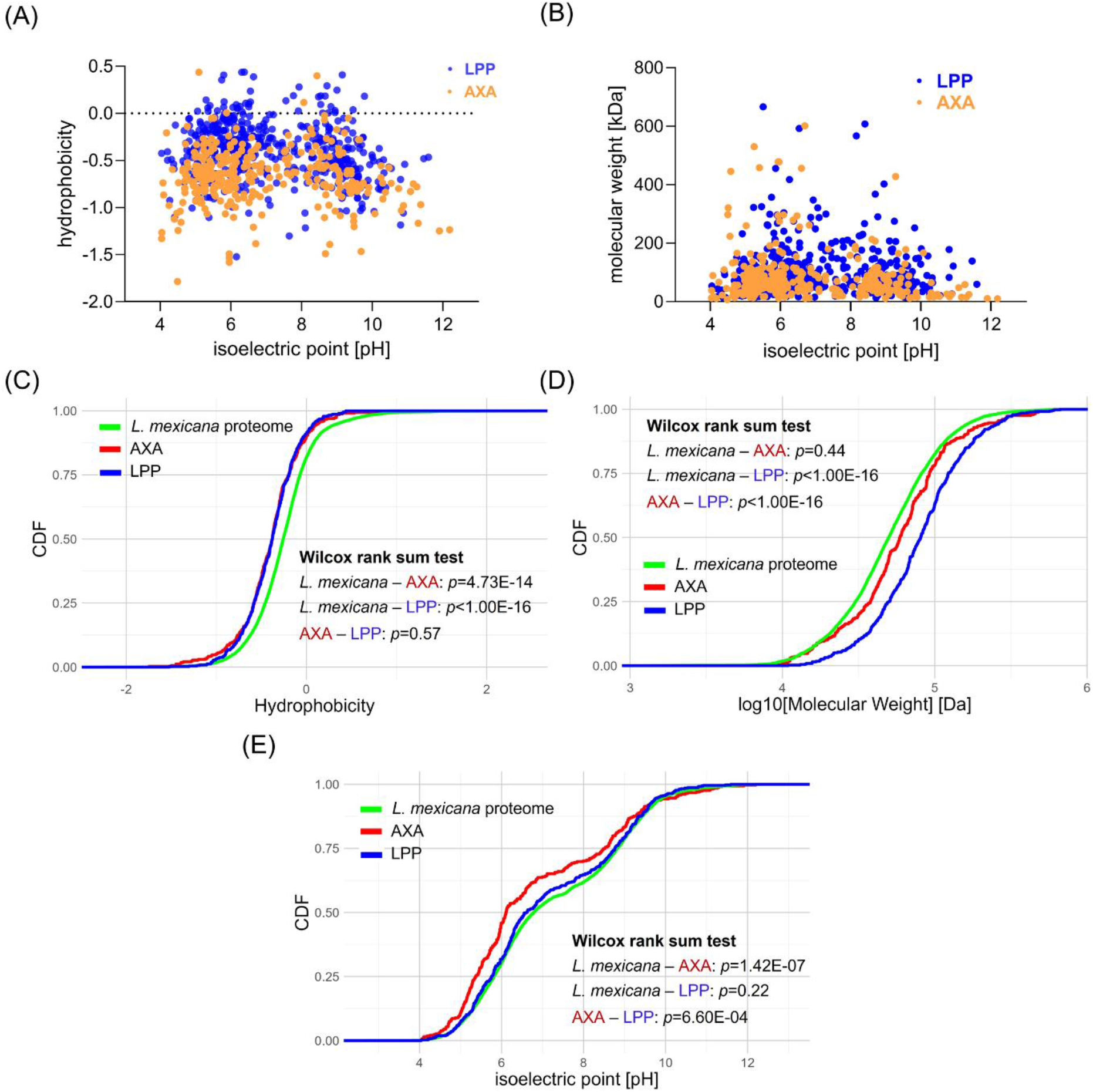
Physicochemical properties of *L. mexicana* phosphorylation substrates. (A) and (B) Scatter plots comparing hydrophobicity and isoelectric points and molecular weights and isoelectric points respectively of phosphorylation substrates in log-phase promastigote (LPP) and axenic amastigote (AXA) life cycle stages. (C), (D) and (E) Cumulative distributions of hydrophobicity, molecular weights and isoelectric points respectively in the AXA and LPP phosphorylation substrates and the entire *L. mexicana* proteome. Wilcox rank sum test *p* values of the comparisons *L. mexicana* total proteome vs AXA phosphorylation substrates (*L. mexicana* – AXA), *L. mexicana* total proteome vs LPP phosphorylation substrates (*L. mexicana* – LPP) and AXA vs LPP phosphorylation substrates (AXA – LPP) are shown.

### HSP90 inhibition differentially affects global protein phosphorylation of *L. mexicana* life cycle stages

Inhibition of HSP90 using tanespimycin caused substantial differences in the global phosphorylation in promastigotes and amastigotes (Fig. 5). Whilst in the LPP, phosphorylation in the majority of PKs were found to decrease with the HSP90 inhibition (Fig. 5A), the opposite trend was observed in the AXA (Fig. 5C). Interestingly, the overall changes in the protein kinome phosphorylation in the SPP was found to lie in between those of the LPP and the AXA (Fig. 5B). Principal Component Analysis (PCA) of the phosphoproteins and the HSP90 inhibition affected phosphoproteins based on their relative quantification profiles revealed clustering of the proteins according to the life cycle stages (Fig. 5D).

**FIG. 5.**
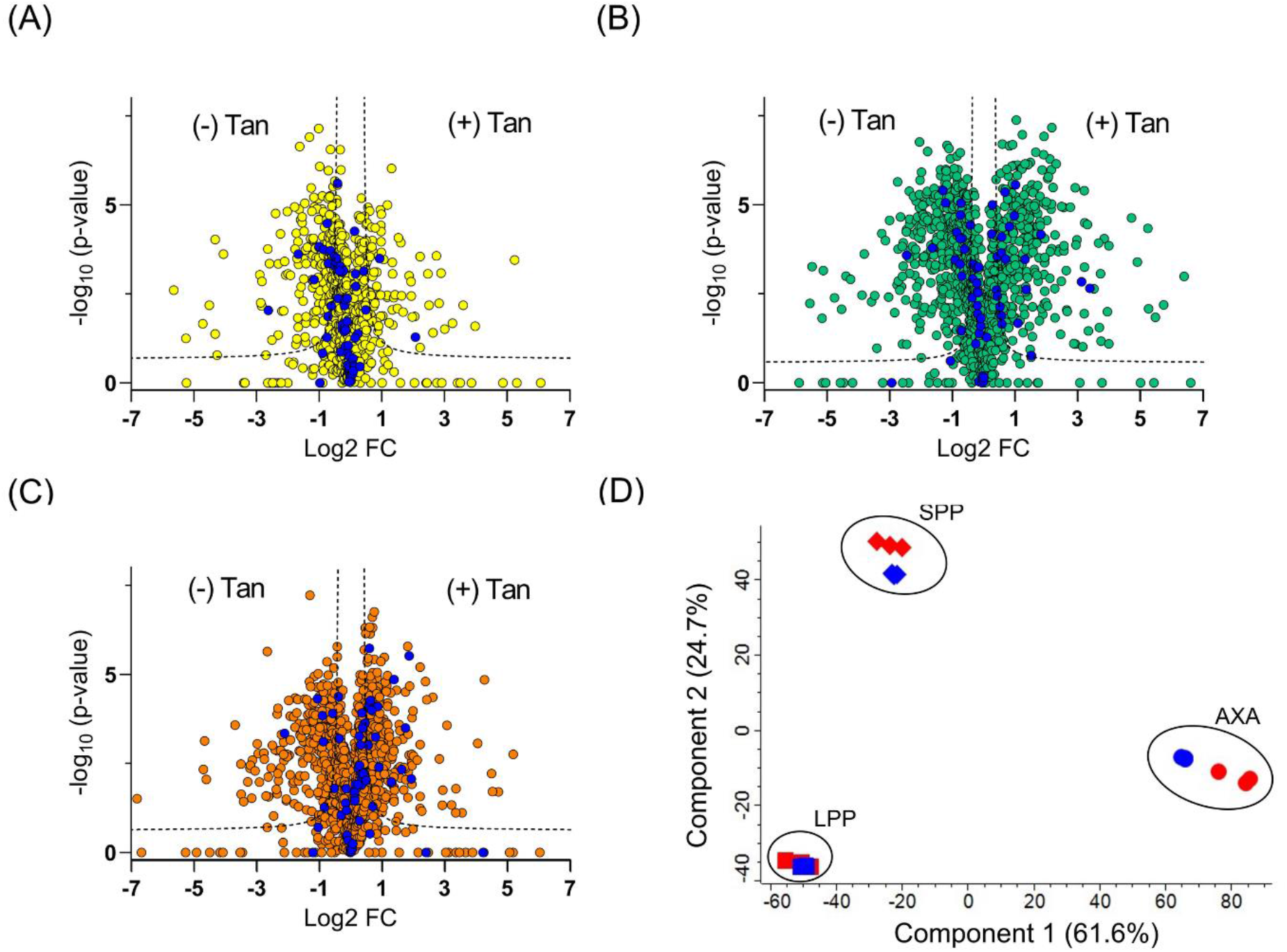
Effect of HSP90 inhibition via tanespimycin treatment on *L. mexicana* phosphoproteome. (A), (B) and (C) Volcano plots showing differential enrichment of phosphoproteins upon tanespimycin treatment (+) Tan and no treatment (-) Tan in log-phase promastigote (LPP), stationary-phase promastigote (SPP) and axenic amastigote (AXA) life cycle stages respectively profiled by phosphoproteome enrichment followed by TMT labelling-based quantitative proteomic MS. All experiments were performed in three biological replicates. A modified *t* test with permutation-based FDR statistics (250 permutations, FDR = 0.05) was applied to compare the quantitative differences in the phosphoproteins between the tanespimycin treated and non-treated groups. Protein kinase are highlighted in blue filled circles. (D) Principal Component Analysis (PCA) of the phosphoproteins (blue) and the HSP90 inhibition affected phosphoproteins (red) in the three life cycle stages based on their relative quantification profiles.

In the LPP, the HSP90 inhibition negatively affected the phosphorylation of 623 proteins (Supplementary Table S5). These phosphoproteins were enriched in the MFs (Supplementary Fig. S8A), microtubule motor activity (*P* value 5.68e-5) and kinase activity (*P* value 9.76e-5). The top BPs (Supplementary Fig. S8B) of these downregulated proteins were microtubule-based movement (*P* value 1.53e-6) and chromatin assembly or disassembly (*P* value 1.39e-5). The most enriched CCs (Supplementary Fig. S8C) of these negatively affected phosphoproteins were cilium (*P* value 4.77e-9) and cytoskeleton (*P* value 3.63e-8). Tanespimycin treatment also led to statistically significant increased phosphorylation of a total of 236 phosphoproteins in the LPP (Table S5). The upregulated phosphoproteins in the LPPs were enriched in the MFs (Supplementary Fig. S8A), anion binding (*P* value 3.31e-6) and ribonucleotide binding (*P* value 6.96e-6). The top BPs (Supplementary Fig. S8B) of these up-regulated phosphoproteins were microtubule-based process (*P* value 9.54e-5) and protein folding (*P* value 3.07e-4). Interestingly, tanespimycin treatment increased the phosphorylation of both HSP90 and HSP70 in the LPP (Supplementary Fig. S9). The most enriched CCs (Fig. S8C) of the positively regulated phosphoproteins were cilium (*P* value 7.73e-6) and eukaryotic translation initiation factor complex 4F (*P* value 6.57e-5).

In the SPP, tanespimycin treatment negatively and positively affected the phosphorylation of 702 and 455 proteins respectively (Table S6). GO analyses of these HSP90 inhibition-affected phosphoproteins in the SPPs showed similar overall trends as that of the affected proteins in the LPP (Supplementary Fig. S10). However, disparity in the cellular processes of the affected proteins were revealed between the promastigotes and the AXA. For example, in contrast to the promastigotes, tanespimycin treatment in the AXA was found to cause a decrease in the phosphorylation of both HSP90 and HSP70 (Supplementary Fig. S9). In the AXA, the HSP90 inhibition negatively and positively affected the phosphorylation of 429 and 614 proteins, respectively (Table S7). The proteins that showed a decreased phosphorylation upon tanespimycin treatment in the AXA were highly enriched in the MFs (Supplementary Fig. S11), structural constituent of ribosome (*P* value 6.28e-16) and unfolded protein binding (*P* value 1.44e-8). Among the most highly enriched BPs of these phosphoproteins were translation (*P* value 8.45e^-16^) and protein folding (*P* value 2.28e-10). The most enriched CC GO terms were cytosolic ribosome (*P* value 5.14e-14) and cytosolic large ribosomal subunit (*P* value 1.90e-12). The AXA proteins that showed an increased phosphorylation upon tanespimycin treatment were highly enriched in the MFs (Supplementary Fig. S11), mRNA binding (*P* value 7.38e-8) and RNA binding (*P* value 4.56e-6). Among the most highly enriched BPs of these proteins were movement of cell or subcellular component (*P* value 1.05e-3) and ribosomal large subunit assembly (*P* value 2.08e-3). The most enriched CC GO terms of these upregulated AXA phosphoproteins were nucleolus (*P* value 6.37e-9) and eukaryotic translation initiation factor complex 4F (*P* value 5.12e-7).

### Correlation between the global phosphoproteome and protein-RNA interactions upon HSP90 inhibition in *L. mexicana*

We have recently demonstrated that HSP90 inhibition causes widespread perturbation of protein-RNA interactions in *L. mexicana* (13). As phosphorylation-dephosphorylation dynamics of RBPs in higher eukaryotes regulate RNA processing and decay in response to various signals (14), we set out to identify the RBPs among the tanespimycin-affected phosphoproteins in both amastigote and promastigote life cycle stages of *L. mexicana*. Because the HSP90 inhibition predominantly downregulated protein-RNA interactions in both life cycle stages of *L. mexicana* (13), we compared the downregulated RBPs in our published data sets with both up and -downregulated phosphoproteins identified in this study.

In the LPP, 169 out of 623 downregulated phosphoproteins and 76 out of 236 upregulated phosphoproteins were found to be downregulated RBPs (Fig. 6A). Similarly, in the AXA, 162 out of 430 downregulated phosphoproteins and 182 out of 614 upregulated phosphoproteins were found to be downregulated RBPs (Fig. 6B). In the LPP, the upregulated (Fig. 6C) and downregulated (Fig. 6D) RNA-binding phosphoproteins showed alpha tubulin and 40S ribosomal protein S1/3 respectively among the most enriched InterPro domains. In contrast, comparison of interprotein domain occurrences in the RNA-binding phosphoproteins of the AXA revealed that RNA helicase and heat shock protein HSP90 were among the most enriched InterPro domains in the upregulated (Fig. 6E) and downregulated (Fig. 6F) phosphoproteins, respectively. The increased phosphorylation detected in RNA helicases such as the ATP-dependent RNA helicase LmxM.28.1310 upon HSP90 inhibition agrees with previous reports suggesting involvement of the HSP90 in RNA metabolism such as RNA splicing, RNA transport and RNA decay (17–19).

**FIG. 6.**
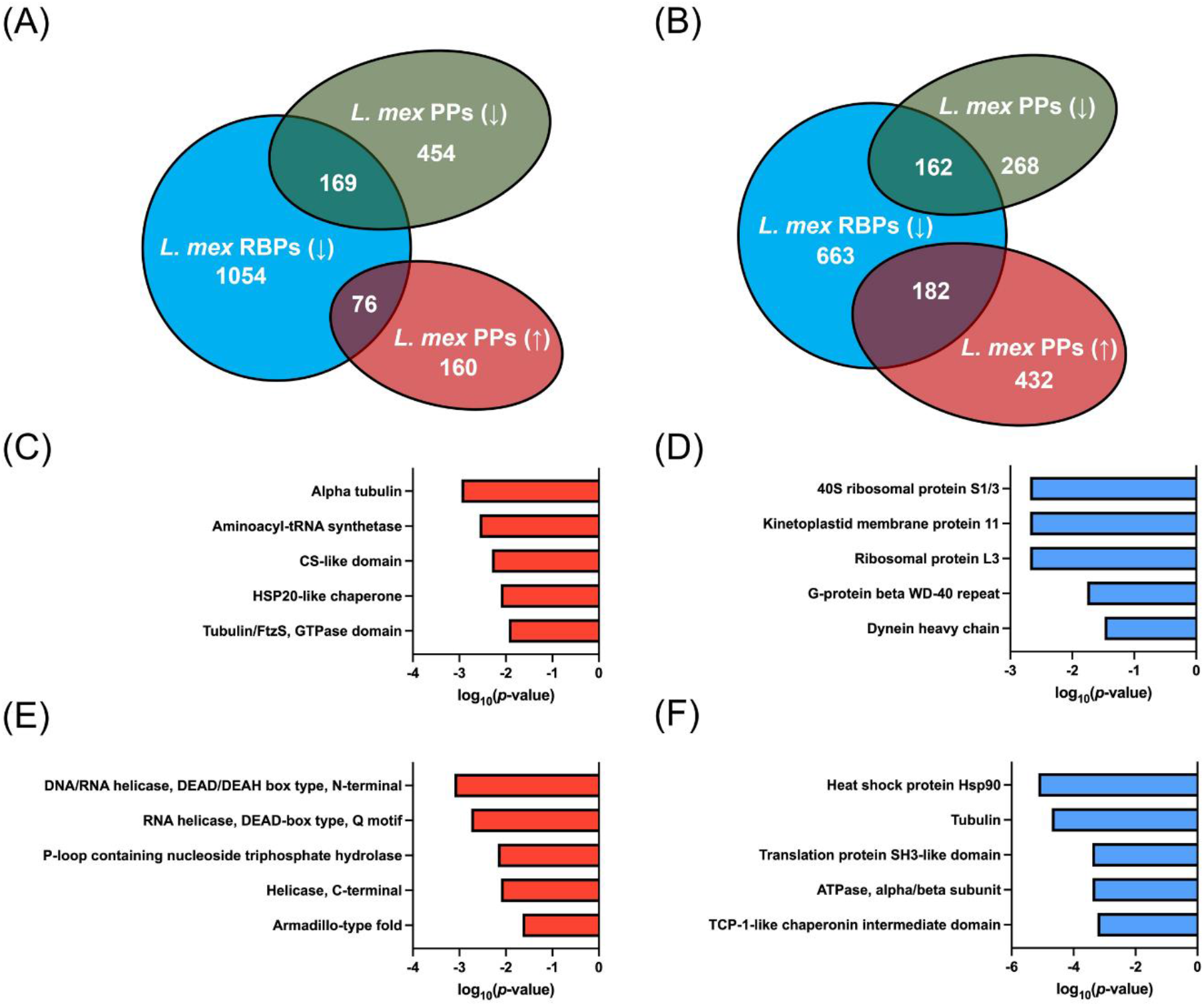
Correlation between RNA-binding proteins (RBPs) downregulated by HSP90 inhibition and the HSP90 inhibition-modulated phosphoproteins in *L. mexicana*. Venn diagrams showing comparison of the downregulated RBPs (*L. mex* RBPs(↓)) and upregulated (*L. mex* PPs(↑)) and downregulated (*L. mex* PPs(↓)) phosphoproteins in (A) log phase promastigote (LPP) and (B) axenic amastigote (AXA) life cycle stages. (C) and (D) most enriched InterPro domains in the upregulated and downregulated *L. mexicana* RNA-binding phosphoproteins respectively in the LPP. (E) and (F) most enriched InterPro domains in the upregulated and downregulated *L. mexicana* RNA-binding phosphoproteins respectively in the AXA.

Next, we constructed protein-protein interaction (PPI) networks of the HSP90 inhibition-perturbed RNA-binding phosphoproteins. Network analysis revealed the important node properties of degree centrality, betweenness centrality and closeness centrality in the four PPI networks (Table S8). In the AXA, the PPI network of the upregulated RNA-binding phosphoproteins revealed cytidine triphosphate synthase, elongation factor Tu, Rac serine-threonine kinase and ATP-dependent DEAD-box helicase as crucial nodes with the highest betweenness centrality (Fig. 7). In contrast, the network of the downregulated AXA phosphoproteins revealed elongation factor-1 gamma and beta tubulin with the highest betweenness centrality (Supplementary Fig. S12). In the LPP downregulated PPI network, actine-like protein and chaperonin alpha subunit were identified with the highest betweenness centrality (Supplementary Fig. S13). In the LPP upregulated PPI network, several nodes were identified with high betweenness centrality (Supplementary Fig. S14). These include glucose-related protein 78, elongation factor 2, nucleolar protein, protein kinase A, stress-induced protein STI1, 14-3-3 protein and chaperonin TCP20. Interestingly, the high betweenness centrality node, Rac serine-threonine kinase, in the AXA upregulated phosphoproteins was identified as a crucial node of information flow in the LPP upregulated phosphoproteins as well, suggesting potential role of phosphorylation of this PK in regulating the HSP90 inhibition stress in both life cycle stages of *Leishmania*. In contrast, the cytidine triphosphate synthase, and the ATP-dependent DEAD-box helicase were unique high betweenness centrality nodes of the AXA phosphoproteome (Fig. 7), suggesting their phosphorylation playing a role in the life cycle-specific regulation of the HSP90 inhibition.

**FIG. 7.**
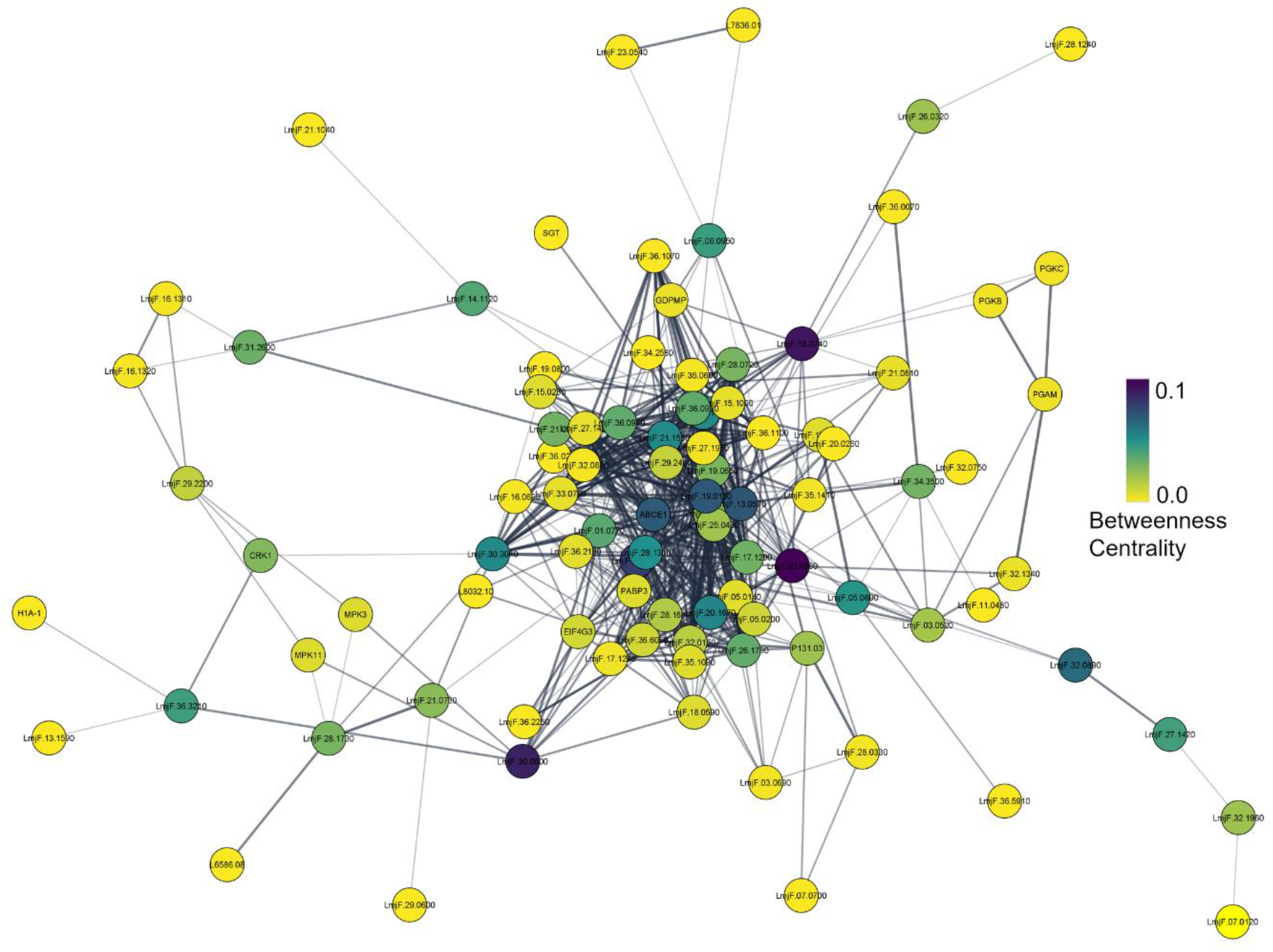
Protein-protein interaction network of RNA-binding phosphoproteins upregulated during HSP90 inhibition in *L. mexicana* axenic amastigotes (AXAs) constructed using publicly available STRING database of *L. major* Friedlin strain. The nodes are coloured according to their betweenness centrality in the network.

## DISCUSSION

The combination of phosphoproteome enrichment with TMT labelling-based quantitative proteomic MS enabled comprehensive profiling of quantitative changes in *L. mexicana* phosphoproteome across its three life cycle stages. Previous studies based on gel-based quantitative methods have shown an increased phosphorylation of HSP90 and its cochaperone HSP70 during the differentiation of *Leishmania* from promastigote to amastigote (6). Our quantitative proteomic MS study not only confirmed these earlier findings but more importantly provided relative changes in the phosphorylation levels of thousands of *Leishmania* proteins during the differentiation. These include many essential PKs, PPs, hydrolases, RBPs, and helicases. We have shown that whilst phosphorylation of chaperone proteins is upregulated in the *L. mexicana* amastigotes, PK phosphorylation was more prominent in the promastigotes. Many PKs are regulated by phosphorylation events (20,21). Phosphorylation of the activation loop of PKs can increase the enzymatic activity by promoting a conformational change that allows the kinase to bind and phosphorylate its substrates more efficiently (20). Conversely, phosphorylation of inhibitory sites can reduce the kinase activity by preventing the activation loop from adopting its catalytically active conformation (20). Thus, the biological effects of phosphorylation of PKs are context dependent and mediated by a network of protein interactions. Our results suggest a higher reliance of the *L. mexicana* promastigotes on the PKs. Nevertheless, our data also accurately captured the upregulation of phosphorylation of a smaller set of PKs in the amastigotes. More importantly, inhibition of HSP90 caused a statistically significant increase in the phosphorylation of many of the amastigote PKs. These include kinases with potential therapeutic targeting scope (22,23) such as the mitogen activated PKs MPK3, MPK10, MPK11, MPK12 and MPK14, cell division PKs CRK1 and CRK3, glycogen synthase kinase GSK-3β and AGC essential kinase-1. It is worth investigating specific targeting of these PKs either alone or in combination with HSP90 inhibition against leishmaniasis.

Phosphorylation can affect the local and global electrostatic potential of substrate proteins (24). It is the location of the phosphorylation site and the specific patterns of residues on both sides of the site within a protein sequence rather than the overall hydrophilicity or hydrophobicity of the protein that makes the protein a substrate of specific kinases in an organism. Interestingly, in *Leishmania*, our results show an overall preference of hydrophilic proteins for phosphorylation in all three life cycle stages. This suggests that *Leishmania* may have evolved a regulatory mechanism for fine-tuning its cellular signalling pathways in response to environmental cues, which can be important for maintaining its cellular homeostasis and adapting to changing conditions.

Protein size and isoelectric point are crucial factors that can influence protein solubility and interactions with other proteins and other biomolecules such as nucleic acids, lipids and metabolites, and can affect the efficiency and specificity of these interactions and signalling pathways (25). Smaller proteins may be more prone to dynamic interactions with other proteins, and proteins with more acidic isoelectric points tend to have a greater propensity for interactions with positively charged proteins and nucleic acids. Therefore, the observed differences in protein size and isoelectric points between phosphorylation substrates in the amastigotes and the promastigotes indicate that there are distinct regulatory networks or signalling pathways that operate at each stage, and these networks may be optimised for different types of protein-protein interactions or cellular responses. It is also possible that these differences reflect changes in the cellular environment or in the expression of specific PKs and PPs that regulate protein phosphorylation. Further research would be needed to determine the specific mechanisms underlying these differences in the phosphorylation of substrate properties between the amastigotes and the promastigotes.

It is conceivable that the changes in the global phosphorylation network caused by the HSP90 inhibition plays a crucial role in the life cycle differentiation of *Leishmania*. Our quantitative proteomic data captured these global changes in the phosphorylation landscape during the transition from the LPP to the SPP to the AXA. It is the amastigote form that causes leishmaniasis in vertebrate hosts. Therefore, identification of druggable targets and pathways in the amastigotes is crucial. The increased phosphorylation of specific PKs detected in the amastigotes following the HSP90 inhibition suggests an increased reliance of the parasite in the amastigote stage on these PKs to mitigate the detrimental effects of the HSP90 inhibition stress. Similarly, the observation that the HSP90 inhibition leads to an increased phosphorylation of RNA helicases in the amastigotes suggests potential role of HSP90 in regulating the phosphorylation status of these important enzymes that unwind and remodel RNA molecules during transcription, translation and RNA decay. It is likely that phosphorylation regulates the enzymatic activity, protein-protein interactions, or localisation of the RNA helicases within the cell. The *L. mexicana* kinesin, LmxM.29.0350, termed DEATH kinesin, was recently found to be essential for the survival of the parasite in the infected host (26). The altered phosphorylation of kinesins detected in the *L. mexicana* upon tanespimycin treatment suggests a crucial role of the HSP90 in regulating cargo transport, transport vesicles, organelles and chromosomes via modulating the phosphorylation of the motor proteins in the parasite. Thus, the findings of this study offer a plethora of valuable insights that can be utilised to explore the molecular mechanisms and implications of inhibiting HSP90 in the *Leishmania* parasite in future research.

## MATERIALS AND METHODS

### Leishmania mexicana culture

*L. mexicana* M379 strain (MNYC/BC/62/M379) promastigotes from frozen stock were quickly defrosted in a water bath at 37°C and inoculated in 10 mL Schneider’s insect medium (Sigma-Aldrich) in T-25 flasks supplemented with 0.4 g/L NaHCO_3_, 0.6 g/L CaCl_2_, and 15% heat-inactivated foetal bovine serum (FBS, Thermo Fisher Scientific, South American origin) at pH 7.0. The parasites were incubated at 26°C for 1-2 days. The parasites were in log phase by this stage with many dividing cells. This culture was used to inoculate another culture by adding 5 x 10^5^ parasites/mL and incubated at 26°C for 2-3 days. In order to generate SPPs, the incubation was continued for 8-9 days. AXAs were generated from the LPPs using changes in pH and temperature of the culture medium as described earlier (13). Briefly, the *L. mexicana* promastigotes in log phase on day 3 of the culture were transferred to 60 mL of pH 5.5 Schneider’s insect medium supplemented with 20% heat-inactivated FBS and incubation continued at 26°C. On day 8-9, the stationary phase parasites were transferred to 60 mL of pH 5.5 Schneider’s insect medium supplemented with 20% heat-inactivated FBS and incubated at 32°C. On day 11-12, the parasites were completely differentiated into amastigote stage. The growth and morphology of parasites were observed under an optical microscope and parasite numbers in cultures at all stages were measured using a haemocytometer.

### Tanespimycin treatment and cell lysate preparation

*L. mexicana* parasites in LPP, SPP and AXA life cycle stages were treated with 1 µM tanespimycin or DMSO (control) in fresh complete Schneider’s insect medium for a total duration of 16 h in biological triplicates. Following treatments, the parasites were washed three times with ice-cold phosphate buffered saline (PBS) and lysed immediately in ice-cold lysis buffer (20 mM Tris. HCl, pH 8.5, 8M urea, phosphatase inhibitors (PhosSTOP, Roche), 2 mM dithiothreitol (DTT) and 1 mM PMSF) by passing the parasites through 29G needles 8-10 times. Lysate debris were eliminated by centrifugation at 16,000 g for 10 min at 4°C. After centrifugation, the protein samples were reduced by DTT treatment (10 mM, 60 min, 35°C), and alkylated by iodoacetamide (IAA, 20 mM, 45 min at room temperature in the dark). The samples were digested overnight at 37°C with sequencing-grade modified trypsin (Promega) at an enzyme to protein ratio of 1:40. The samples were then acidified with trifluoroacetic acid (TFA) (0.1% v/v final concentration; Sigma-Aldrich), centrifuged at 16,000 g for 10 min and the supernatant was collected. The tryptic peptides were then desalted on C-18 Sep-Pak Classic cartridges (Waters; WAT051910) following manufacturer’s instructions. The peptides were evaporated to complete dryness in a speed vacuum concentrator and stored at -80°C until required.

### Phosphopeptide enrichment

Phosphopeptides were enriched using High-Select Fe-NTA Phosphopeptide Enrichment Kit (Thermo Scientific) following manufacturer’s protocols. The eluted phosphopeptides were dried immediately in a speed vacuum concentrator. The samples were then acidified with 0.1% TFA and desalted on C-18 Sep-Pak Classic cartridges (Waters; WAT051910) following manufacturer’s instructions. The peptides were evaporated to complete dryness in a speed vacuum concentrator and subjected to TMT labelling.

### TMT labelling

TMT labelling of the desalted phosphopeptides was carried out using TMTsixplex isobaric label reagent set (Thermo Fisher Scientific). The phosphopeptides of each experimental condition were dissolved in 100 µL of 100 mM triethylammonium bicarbonate (TEAB) and treated with room temperature equilibrated and freshly dissolved unique TMT label reagent in 41 µL anhydrous acetonitrile. The labelling reactions were run for 1 h at room temperature, following which 10 µL of 5% solution of hydroxylamine was added and incubated for 15 min at room temperature to quench the reactions. The six samples of three replicates of each life cycle stage were then combined together and concentrated to complete dryness in a speed vacuum concentrator. The samples were then redissolved in 0.1% TFA, desalted and cleaned-up using Pierce Peptide Desalting Spin Columns (Thermo Fisher Scientific) following manufacturer’s instructions. The samples were dried in a speed vacuum concentrator and stored at -80 °C until required.

### Nano LC-MS/MS data acquisition

The LC-MS/MS analyses of TMT-labelled peptides were performed on an Orbitrap Fusion Lumos Mass Spectrometer (Thermo Fisher Scientific) coupled with a Thermo Scientific Ultimate 3000 RSLCnano UHPLC system (Thermo Fisher Scientific). Desalted and TMT-labelled tryptic peptides dissolved in 0.1% formic acid (FA) were first loaded onto an Acclaim PepMap 100 C18 trap column (5 µm particle size, 100 µm id X 20 mm, TF164564) heated to 45 °C using 0.1% FA/H_2_O with a flow rate of 10 µL/min, then separated on an Acclaim PepMap 100 NanoViper C18 column (2 µm particle size, 75 µm id X 50 cm, TF164942) with a 5% to 38% ACN gradient in 0.1% FA over 125 min at a flow rate of 300 nL/min. The full MS spectra (*m/z* 375 to 1,500) were acquired in Orbitrap at 120,000 resolution with an AGC target value of 4e^5^ for a maximum injection time of 50 ms. High-resolution HCD MS2 spectra were generated in positive ion mode using a normalised collision energy of 38% within a 0.7 *m/z* isolation window using quadrupole isolation. The AGC target value was set to 10e^4^, and the dynamic exclusion was set to 45 s. The MS2 spectra were acquired in Orbitrap with a maximum injection time of 54 ms at a resolution of 30,000 with an instrument determined scan range beginning at *m/z* 100. To ensure quality peptide fragmentation a number of filters were utilised, including peptide monoisotopic precursor selection, minimum intensity exclusion of 10e^3^ and exclusion of precursor ions with unassigned charge state as well as charge state of +1 or superior to +7 from fragmentation selection. To prevent repeat sampling, a dynamic exclusion with exclusion count of 1, exclusion duration of 30 s, mass tolerance window of +/-7 ppm and isotope exclusion were used.

### Proteome data processing and analysis

All raw LC-MS/MS data were processed using MaxQuant software (27) version 1.6.3.4 with integrated Andromeda database search engine (28). The MS/MS spectra were queried against *L. mexicana* sequences from UniProt KB (8,559 sequences). The following search parameters were used: reporter ion MS2 with multiplicity 6plex TMT, trypsin digestion with maximum 2 missed cleavages, carbamidomethylation of cysteine as a fixed modification, oxidation of methionine, acetylation of protein N-termini and phosphorylation of serine, threonine and tyrosine residues as variable modifications, minimum peptide length of 6, maximum number of modifications per peptide set at 5, and protein false discovery rate (FDR) 0.01. Appropriate correction factors for the individual TMT channels for both lysine side-chain labelling and peptide N-terminal labelling as per the TMT-6plex kits used (Thermo Fisher Scientific) were configured into the database search. The proteinGroups.txt files from the MaxQuant search outputs were processed using Perseus software (29) version 1.6.2.3. Sequences only identified by site, reverse sequences, and potential contaminants were filtered out. A requirement of six non-zero valid value were set across the eighteen reporter intensity corrected main columns of the three life cycle stages. The reporter intensities were normalised by Z-score and transformed to log2 scale. Proteins identified with fewer than 2 unique peptides were discarded and a modified *t* test with permutation-based FDR statistics (250 permutations) was applied to compare the different life cycle stages and tanespimycin-treated and non-treated groups.

### Bioinformatic analysis

Gene Ontology (GO) Terms (Molecular Function, Biological Process, and Cellular Component) of the phosphoprotein data sets were derived from TriTrypDB (tritrypdb.org) (30). Custom R scripts with R 64 bit version 4.2.3 along with R package ggplot2 (version 3.4.2) were used for visualising the GO Terms. InterPro domain occurrences in the phosphoproteins were derived from bioinformatics analysis of the protein IDs using the Database for Annotation, Visualization and Integrated Discovery (DAVID) bioinformatics resources version 6.8 (31). Grand average hydrophobicity values of the phosphoproteins and the entire *L. mexicana* proteome were calculated using the ExPASy (32) tool ProtParam. Isoelectric points and molecular weights were computed using the ExPASy tool Compute pI/Mw. Cumulative distributions of physicochemical properties were derived using custom R scripts and visualised using the R package ggplot2. Protein-protein interaction network analyses were performed by using the publicly available STRING database (version 11.5) (33) of *L. major* strain Friedlin. The open source software platform Cytoscape (version 3.9.1) (34) was used for refining, analysing, and visualising the protein interaction network.

### Data availability

All raw mass spectrometry proteomics data have been deposited to the ProteomeXchange Consortium via the PRIDE partner repository with the dataset identifiers PXD043002.

## SUPPLEMENTAL MATERIAL

**FIG. S1**, PDF file

**FIG. S2**, PDF file

**FIG. S3**, TIF file

**FIG. S4**, TIF file

**FIG. S5**, TIF file

**FIG. S6**, TIF file

**FIG. S7**, TIF file

**FIG. S8**, TIF file

**FIG. S9**, TIF file

**FIG. S10**, TIF file

**FIG. S11**, TIF file

**FIG. S12**, TIF file

**FIG. S13**, TIF file

**FIG. S14**, TIF file

**TABLE S1**, XLSX file

**TABLE S2**, XLSX file

**TABLE S3**, XLSX file

**TABLE S4**, XLSX file

**TABLE S5**, XLSX file

**TABLE S6**, XLSX file

**TABLE S7**, XLSX file

**TABLE S8**, XLSX file

## ACKNOWLEDGEMENTS

This work of E. O. J. P, P. W. D and P. G. S was supported by funding from MRC-Global Challenges Research Fund-Neglected Tropical Diseases (Grant number: MR/P027989/1A). L. G thank funding from the National Natural Science Foundation of China (Grant number: U1801681) and the National Key R&D Program of China (Grant number: 2022YFA1104800). The work of K. K was funded by the MRC-Global Challenges Research Fund-Neglected Tropical Diseases (Grant number: MR/P027989/1A), The Royal Society Research Grant: (Grant number: RGS\R2\222343) and the Royal Society of Chemistry Research Fund (Grant number: R21-3545544506).

## SUPPLEMENTAL MATERIAL LEGEND

**FIG. S1** MS/MS spectra of phosphorylated tryptic peptides from L. mexicana HSP90.

**FIG. S2** MS/MS spectra of phosphorylated tryptic peptides from selected L. mexicana proteins.

**FIG. S3** AXA vs LPP with Pfam annotation bar chart.

**FIG. S4** AXA vs LPP (A) and AXA vs SPP (B) volcano plots with kinases highlighted in blue.

**FIG. S5** Biological Process (BP) Gene Ontology (GO) terms enriched in (A) LPP, (B) SPP and (C) AXA phosphoproteins.

**FIG. S6** Cellular Component (CC) Gene Ontology (GO) terms enriched in (A) LPP, (B) SPP and (C) AXA phosphoproteins.

**FIG. S7** Physicochemical properties of *L. mexicana* phosphorylation substrates. (A) and (B) Scatter plots comparing hydrophobicity and isoelectric points and molecular weights and isoelectric points respectively of phosphorylation substrates in stationary phase promastigote (SPP) and axenic amastigote (AXA) life cycle stages. (C), (D) and (E) Cumulative distributions of hydrophobicity, molecular weights and isoelectric points respectively in the AXA and SPP phosphorylation substrates and the entire *L. mexicana* proteome. Wilcox rank sum test *p* values of the comparisons *L. mexicana* total proteome vs AXA phosphorylation substrates (*L. mexicana* – AXA), *L. mexicana* total proteome vs SPP phosphorylation substrates (*L. mexicana* – SPP) and AXA vs SPP phosphorylation substrates (AXA – SPP) are shown.

**FIG. S8** Gene Ontology (GO) analysis of tanespimycin-affected phosphoproteins in *L. mexicana* log phase promastigotes (LPP). (A), (B) and (C) are Molecular Function (MF), Biological Process (BP) and Cellular Component (CC) GO terms respectively; blue: negatively affected phosphoproteins, red: positively affected phosphoproteins.

**FIG. S9** Profile plot of the *L. mexicana* phosphoproteins with (+) and without (-) tanespimycin (tan) treatment in axenic amastigote (AXA), log phase promastigote (LPP) and stationary phase promastigote (SPP) life cycle stages. HSP90 (blue) and HSP70 (red) proteins are highlighted.

**FIG. S10** Gene Ontology (GO) analysis of tanespimycin-affected phosphoproteins in *L. mexicana* stationary phase promastigotes (SPP). (A), (B) and (C) are Molecular Function (MF), Biological Process (BP) and Cellular Component (CC) GO terms respectively; blue: negatively affected phosphoproteins, red: positively affected phosphoproteins.

**FIG. S11** Gene Ontology (GO) analysis of tanespimycin-affected phosphoproteins in *L. mexicana* axenic amastigotes (AXA). (A), (B) and (C) are Molecular Function (MF), Biological Process (BP) and Cellular Component (CC) GO terms respectively; blue: negatively affected phosphoproteins, red: positively affected phosphoproteins.

**FIG. S12** Protein-protein interaction network of RNA-binding phosphoproteins downregulated during HSP90 inhibition in *L. mexicana* axenic amastigotes (AXAs) constructed using publicly available STRING database of *L. major* Friedlin strain. The nodes are coloured according to their betweenness centrality in the network.

**FIG. S13** Protein-protein interaction network of RNA-binding phosphoproteins downregulated during HSP90 inhibition in *L. mexicana* log phase promastigotes (LPPs) constructed using publicly available STRING database of *L. major* Friedlin strain. The nodes are coloured according to their betweenness centrality in the network.

**FIG. S14** Protein-protein interaction network of RNA-binding phosphoproteins upregulated during HSP90 inhibition in *L. mexicana* log phase promastigotes (LPPs) constructed using publicly available STRING database of *L. major* Friedlin strain. The nodes are coloured according to their betweenness centrality in the network.

