## Supplementary Information for "Inhibition of HSP90 distinctively modulates the global phosphoproteome of *Leishmania mexicana* developmental stages"

MTETFAFQAEINQLMSLIINTFYSNKEIFLRELISNA**S**$_{38}$DACDKIRYQ**S**$_{48}$LTDPSVLGDATRLCVRVVPDKENKTLTV
EDNGIGMTKADLVNNLGTIARSGTKAFMEALEAGGDMSMIGQFGVGFYSAYLVADRVTVTSKNNSDEVYVWESS
AGGTFTITSAPESDMKRGTRITLHLKEDQLEYLEVRRLKELIKKHSEFIGYDIELMVEKT**T**$_{211}$EKEV**T**$_{216}$DEDEEEAKK
ADEDGEEPKVEEV**T**$_{239}$EGEEGKKKKTKKVKEV**T**$_{256}$KEYEVQNKHKPLWTRDPKDVTKEEYAAFYKAI**S**$_{289}$NDWE
DPAATKHFSVEGQLEFRSIMFVPKRAPFDMFEPNKKRNNIKLYVRRVFIMDNCEDLCPDWLGFVKGVVDSEDLPLN
I**S**$_{371}$RENLQQNKILKVIRKNIVKKCLEMFEEVAENKEDYKQFYEQFGKNIKLGIHEDTANRKKLMELLRFYSTESGEE
MTTLKDYVTRMKAEQKSIYYITGDSKKKLES**S**$_{477}$PFIEQAKRRGFEVLFMTEPIDEYVMQQVKDFEDKKFACLTKEG
VHFEE**S**$_{526}$EEEKRQREEEKAACEKLCKTMKEVLGDKVEKVTVSERLSTSPCILVTSEFGWSAHMEQIMRNQALRD**S**
$_{594}$**S**$_{595}$MAQYMMSKKTMELNPKHPIIKELRRRVEADENDKAVKDLVFLLFDTSLLTSGFQLEDPTGYAERINRMIKLG
LSLDEEEEEAVEAAVAETAPAEVTAGTSSMEQVD

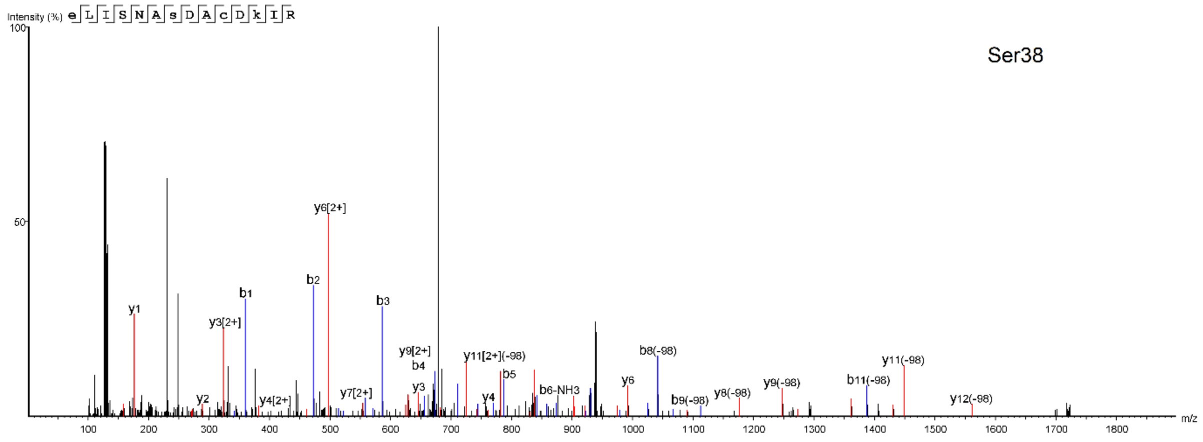


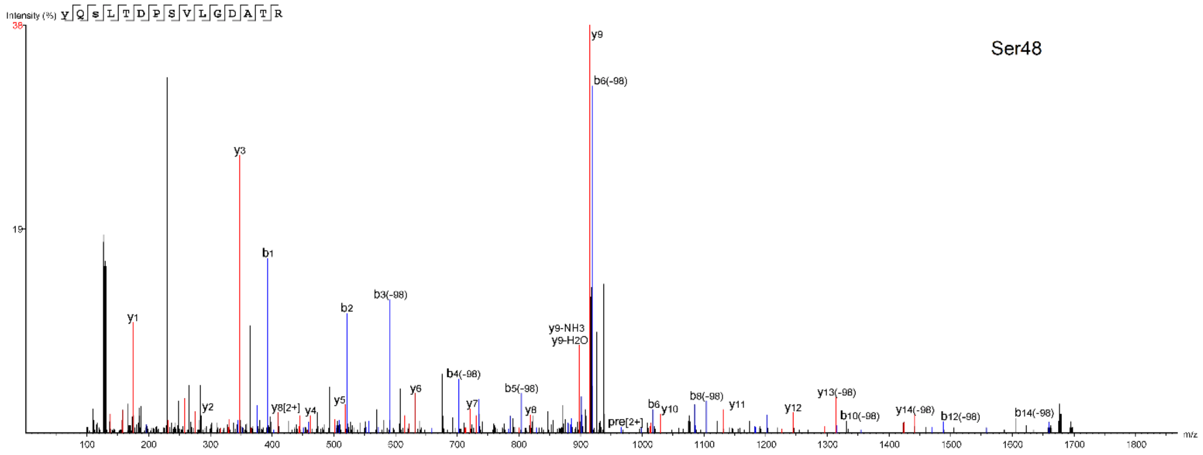

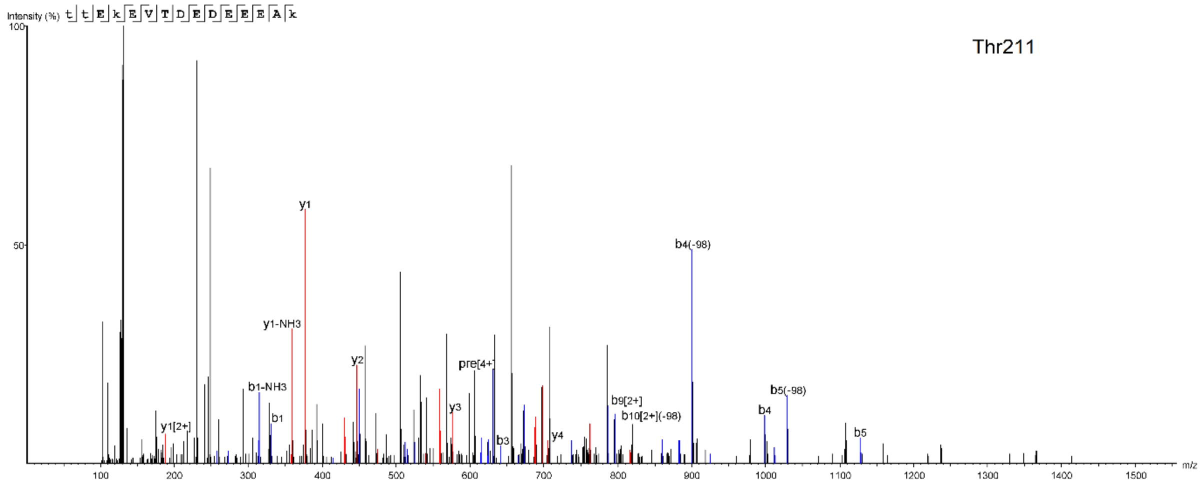
Thr211
Intensity (%)
t t E k E V T D E D E E E A k
y1
y1-NH3
y2
y3
y4
y1[2+]
b1-NH3
b1
b3
b4
b5
b4(-98)
b5(-98)
b9[2+]
b10[2+](-98)
pre[4+]
m/z

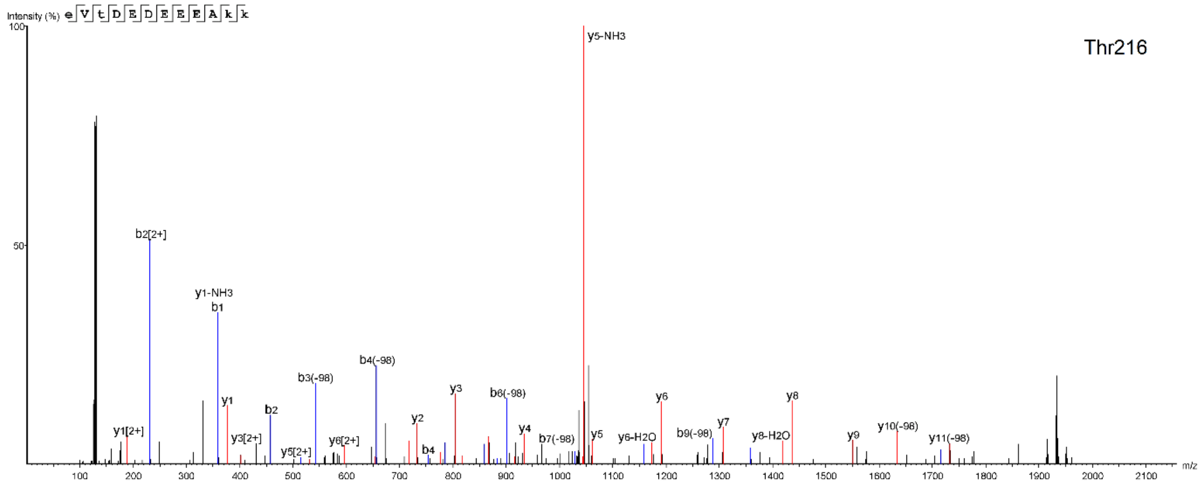
Thr216
Intensity (%)
e V t D E D E E E A k k
y5-NH3
b2[2+]
y1-NH3
b1
y1
y1[2+]
y3[2+]
y5[2+]
y6[2+]
b2
b3(-98)
b4(-98)
b4
y2
y3
b6(-98)
y4
b7(-98)
y5
y6-H2O
y6
b9(-98)
y7
y8-H2O
y8
y9
y10(-98)
y11(-98)
m/z

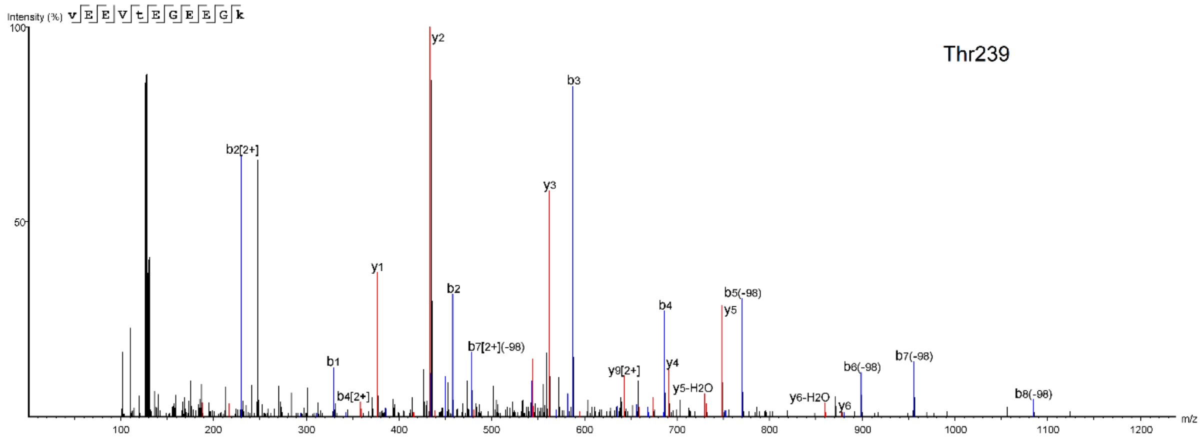
Thr239
Intensity (%)
v E E V t E G E E G k
y2
b3
b2[2+]
y3
y1
b2
b1
b4[2+]
b7[2+](-98)
y9[2+]
b4
y4
y5-H2O
b5(-98)
y5
y6-H2O
y6
b6(-98)
b7(-98)
b8(-98)
m/z

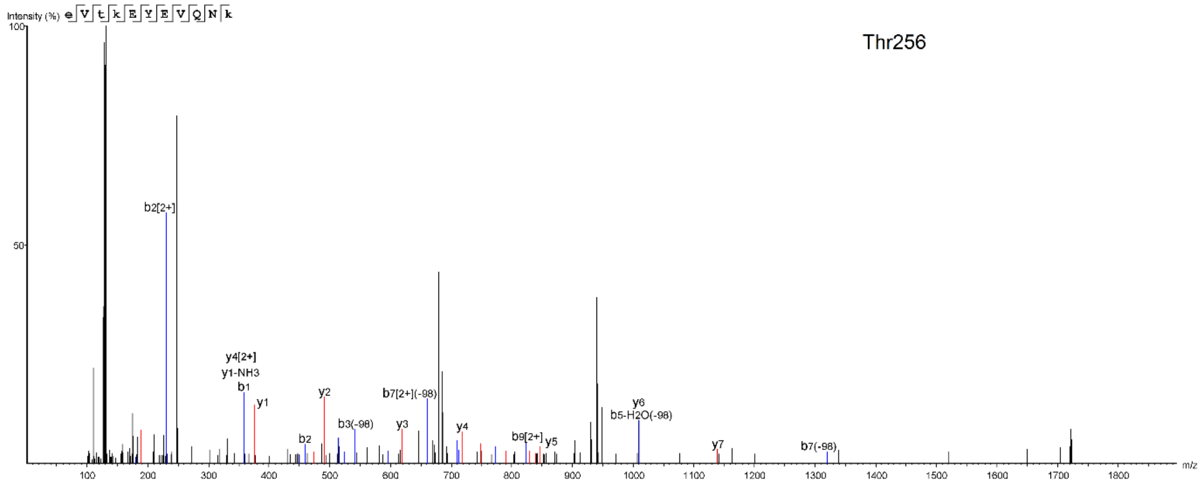
Thr256
Intensity (%)
e V t k E Y E V Q N k
b2[2+]
y4[2+]
y1-NH3
b1
y1
y2
b2
b3(-98)
y3
b7[2+](-98)
y4
b9[2+]
y5
y6
b5-H2O(-98)
y7
b7(-98)
m/z

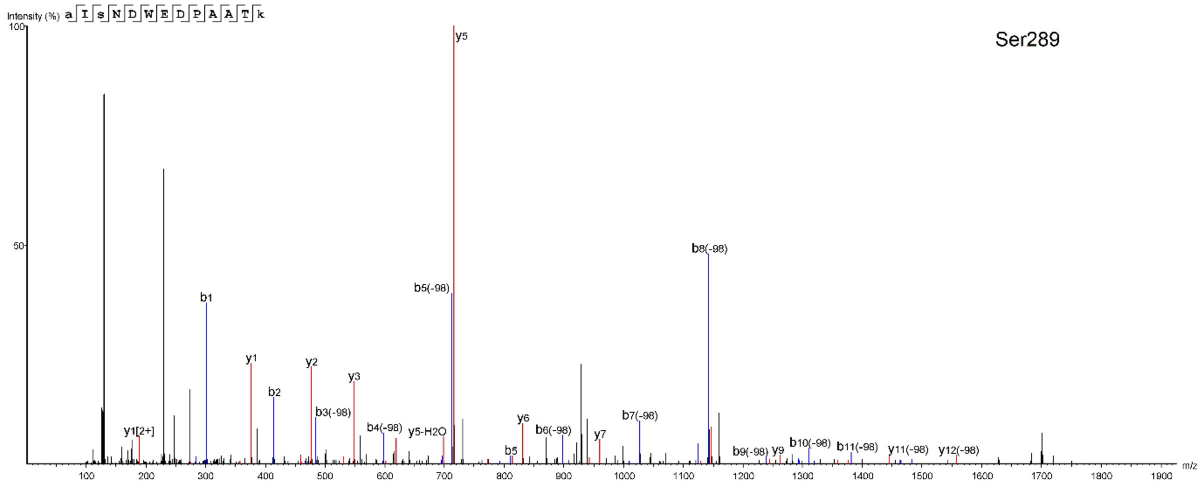
Ser289
Intensity (%)
a I s N D W E D P A A T k
y5
b8(-98)
b5(-98)
b1
y1
y2
y3
b2
b3(-98)
y1[2+]
b4(-98)
y5-H2O
b5
y6
b6(-98)
y7
b7(-98)
b9(-98)
y9
b10(-98)
b11(-98)
y11(-98)
y12(-98)
m/z

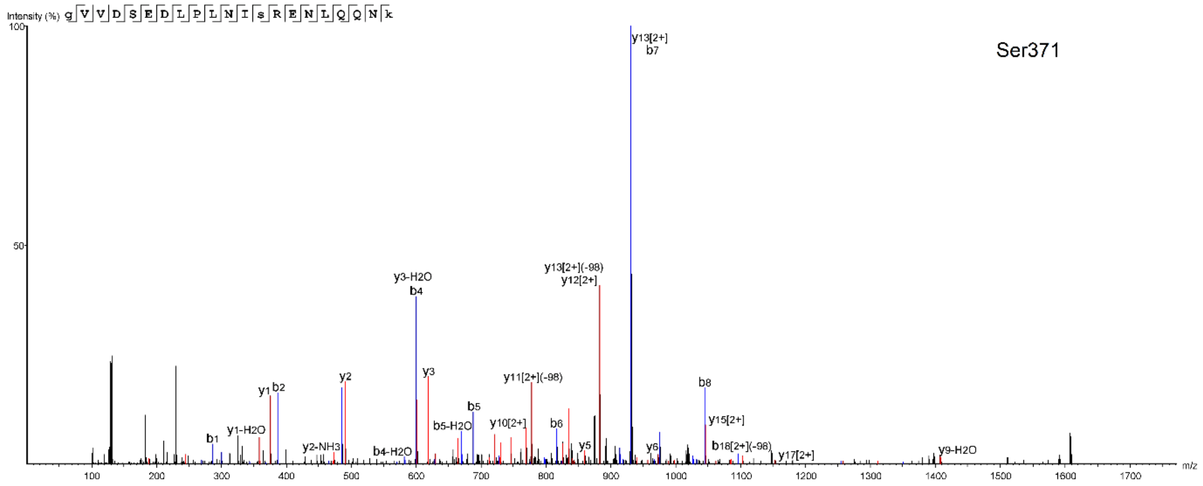
Ser371
Intensity (%)
g V V D S E D L P L N I s R E N L Q Q N k
y13[2+]
b7
y13[2+](-98)
y12[2+]
y3-H2O
b4
y3
y2
y1
b2
y11[2+](-98)
b8
b5
b5-H2O
y10[2+]
b6
y15[2+]
b1
y1-H2O
y2-NH3
b4-H2O
y5
y6
b18[2+](-98)
y17[2+]
y9-H2O
m/z

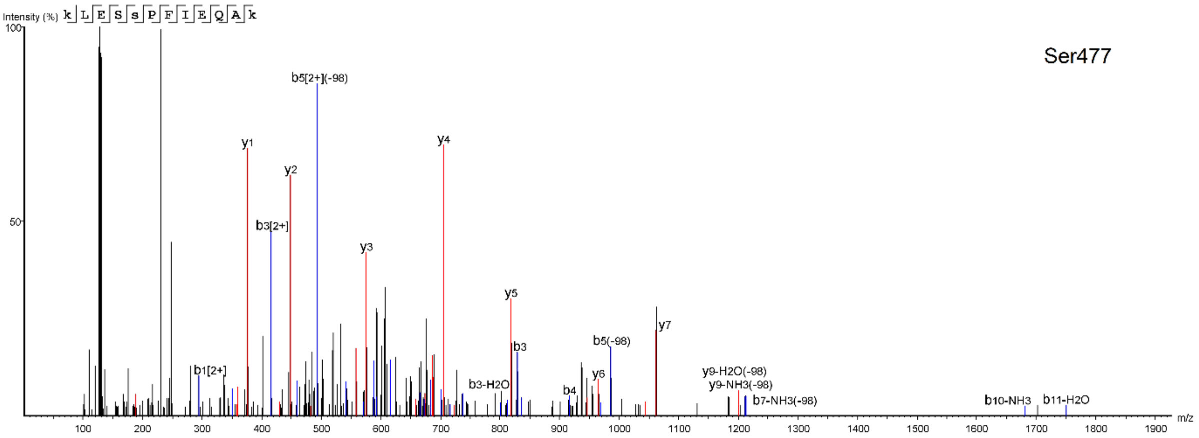
Intensity (%)
k L E S s P F I E Q A k
Ser477
b5[2+](-98)
y1
y4
y2
b3[2+]
y3
y5
y7
b3
b5(-98)
b1[2+]
y6
y9-H2O(-98)
y9-NH3(-98)
b3-H2O
b4
b7-NH3(-98)
b10-NH3
b11-H2O
m/z

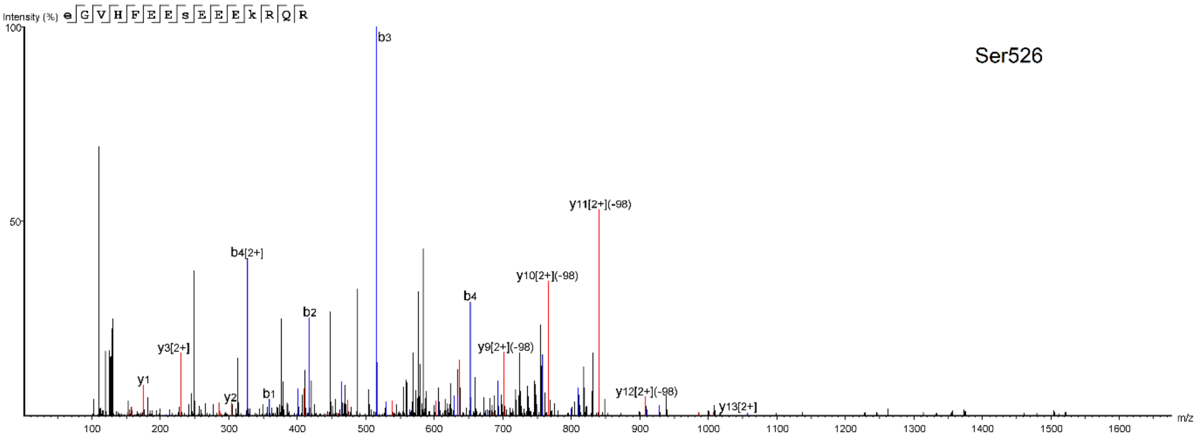
Intensity (%)
e G V H F E E s E E E k R Q R
Ser526
b3
y11[2+](-98)
b4[2+]
y10[2+](-98)
b4
b2
y3[2+]
y9[2+](-98)
y1
b1
y2
y12[2+](-98)
y13[2+]
m/z

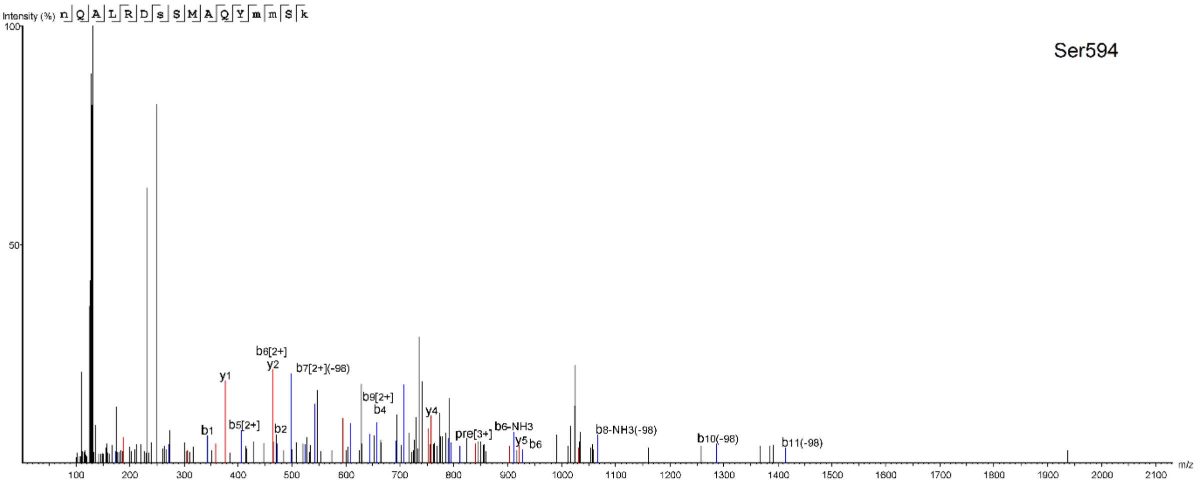
Intensity (%)
n Q A L R D s S M A Q Y m m S k
Ser594
b6[2+]
y2
b7[2+](-98)
y1
b9[2+]
b4
y4
b1
b5[2+]
b2
pre[3+]
b6-NH3
y5
b6
b8-NH3(-98)
b10(-98)
b11(-98)
m/z

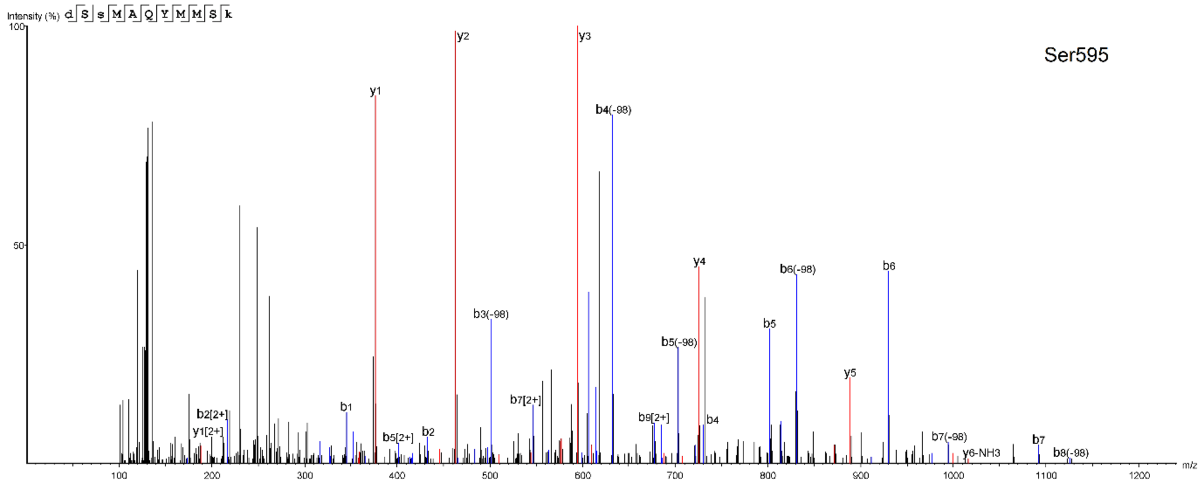


**FIG. S1** MS/MS spectra of phosphorylated tryptic peptides from *L. mexicana* HSP90

Mitogen-activated protein kinase 2 (MPK2)

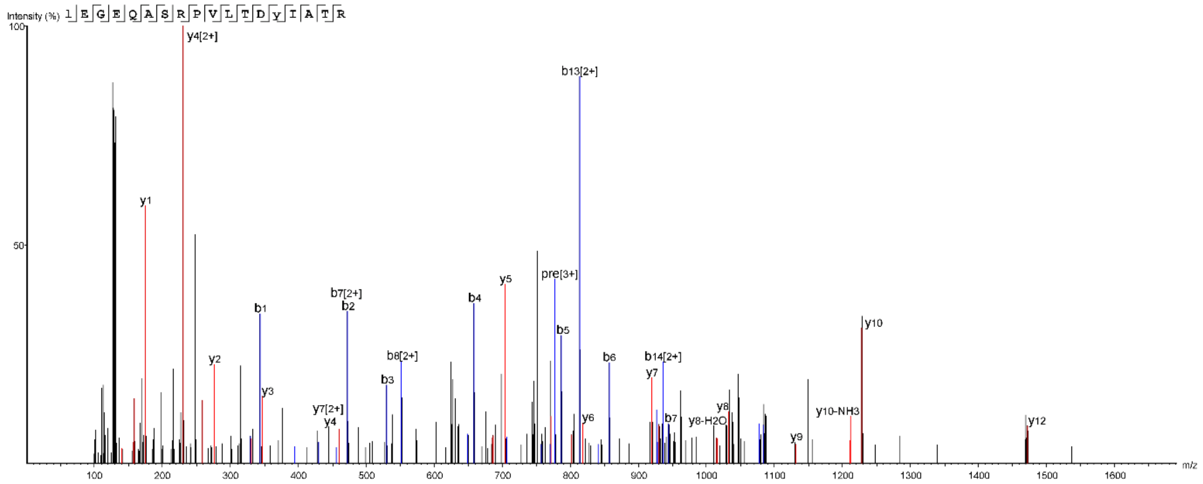


Mitogen-activated protein kinase 3 (MPK3)

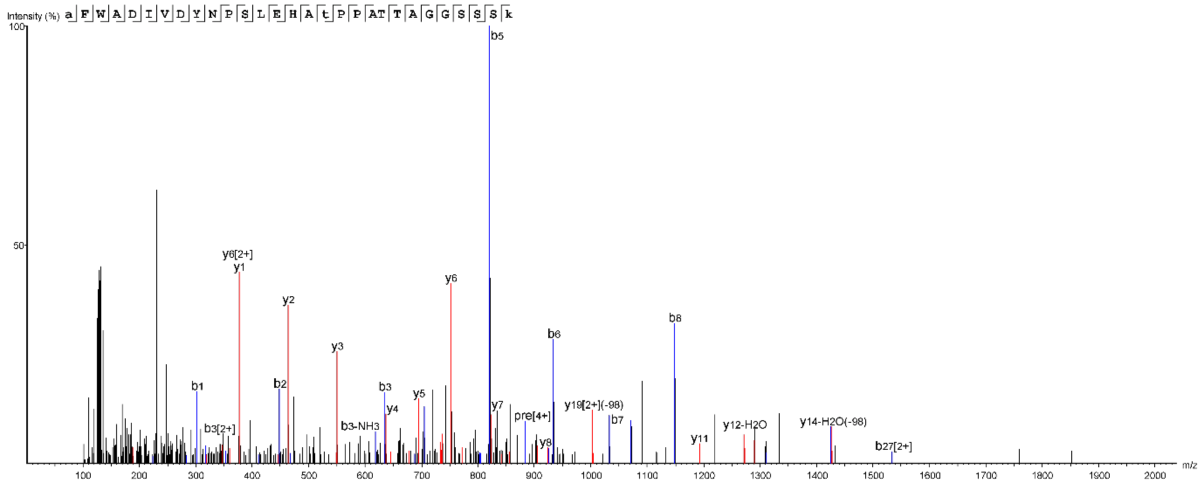

### Mitogen-activated protein kinase 5 (MPK5)

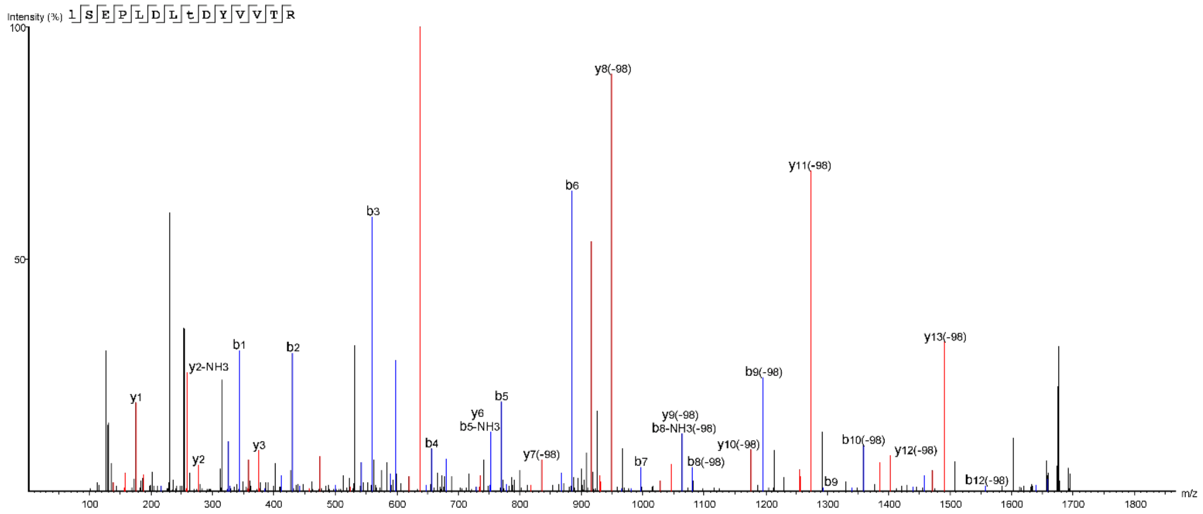


### Mitogen-activated protein kinase 6 (MPK6)

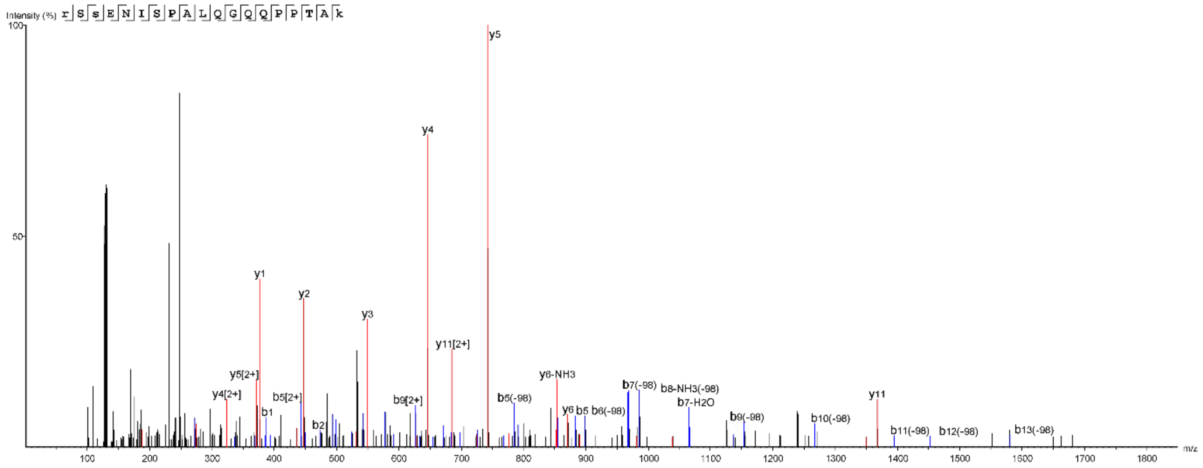


### Mitogen-activated protein kinase 7 (MPK7)

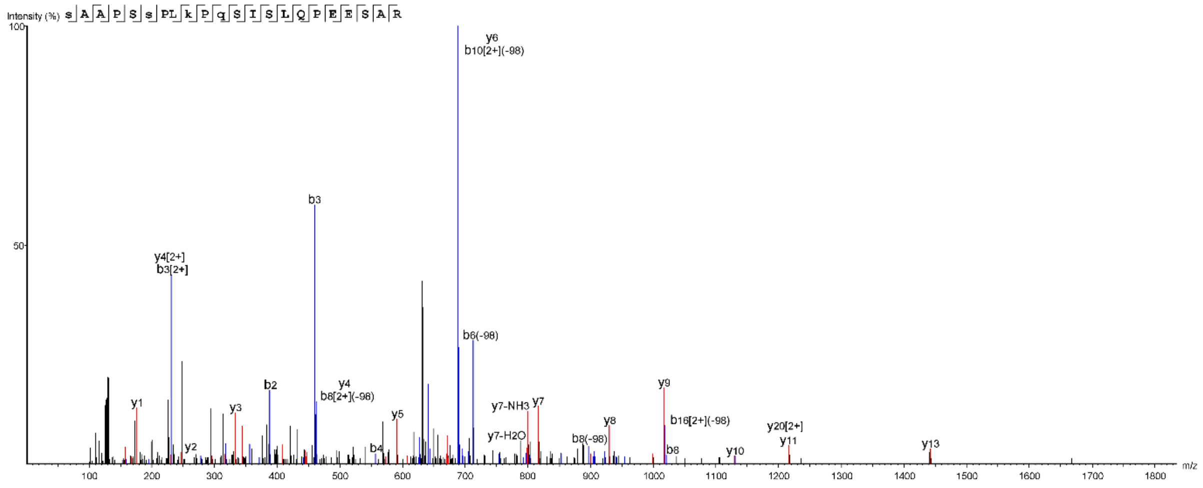

### Mitogen-activated protein kinase 9 (MPK9)

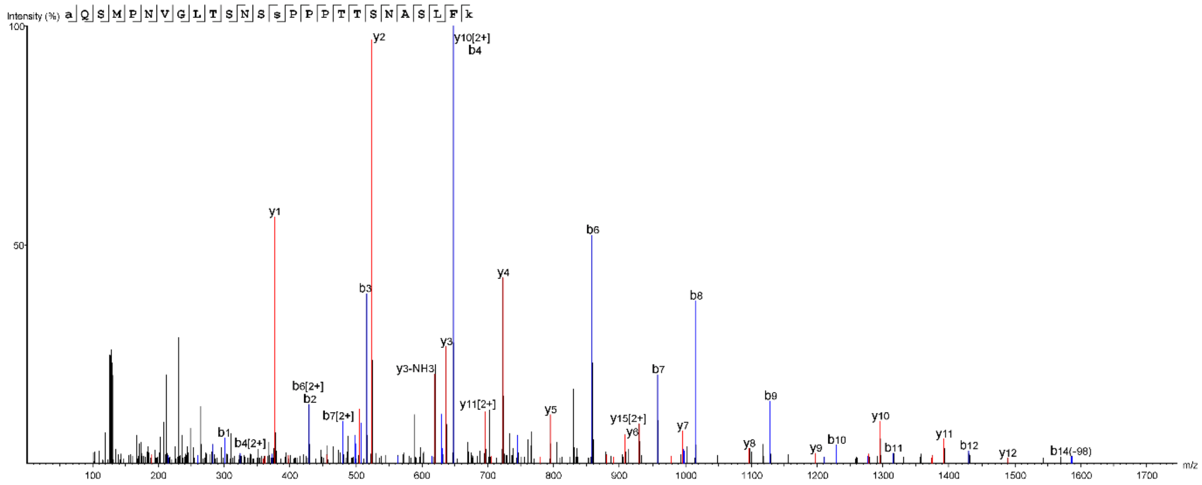


### Mitogen-activated protein kinase 11 (MPK11)

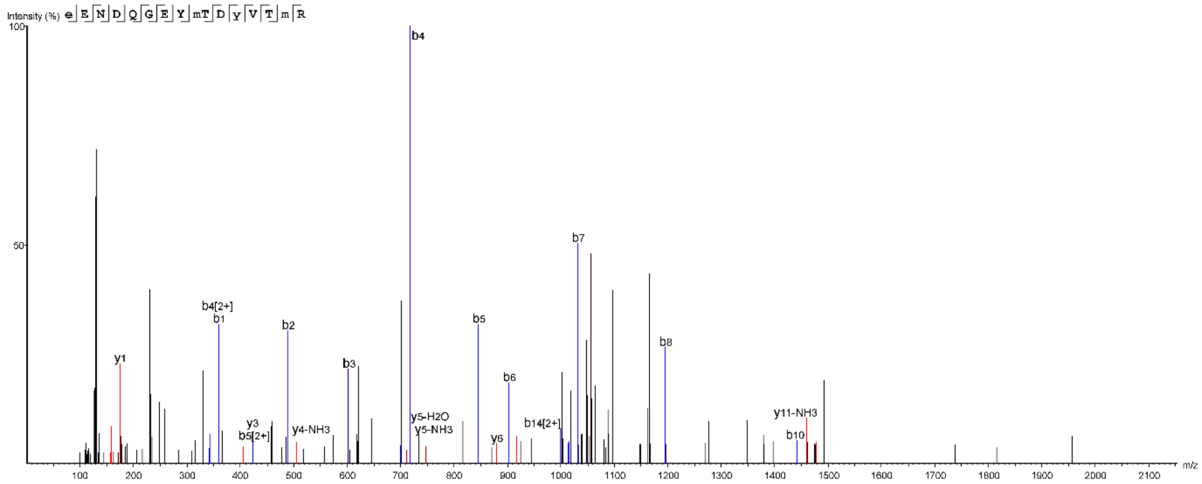


### Mitogen-activated protein kinase 12 (MPK12)

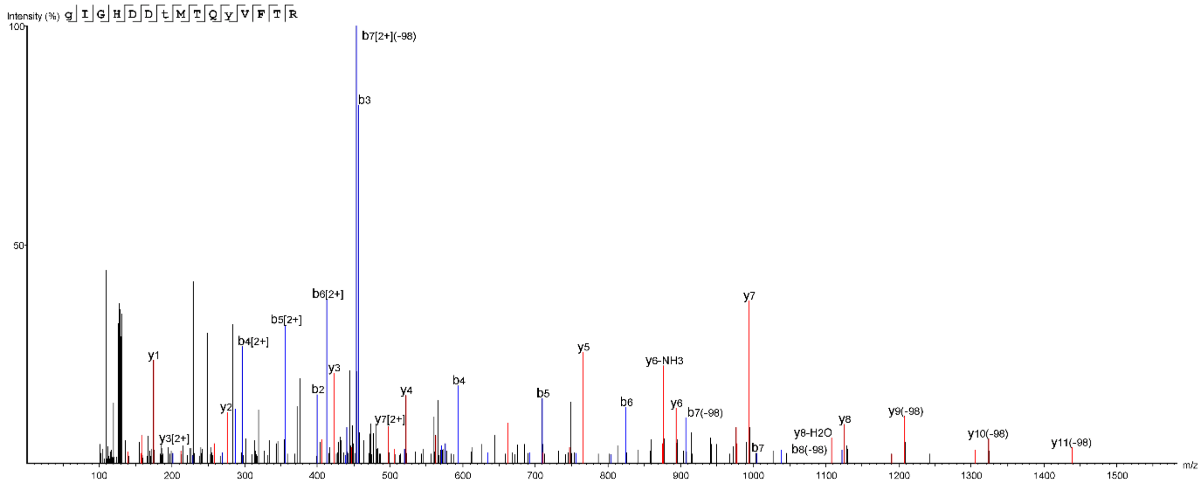

### Mitogen-activated protein kinase 14 (MPK14)

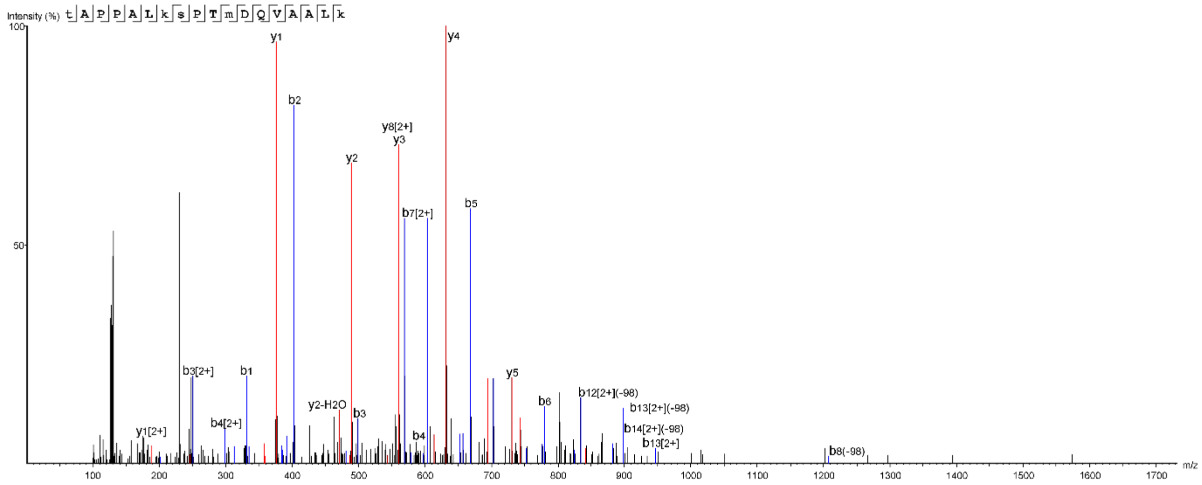


### Mitogen-activated protein kinase 15 (MPK15)

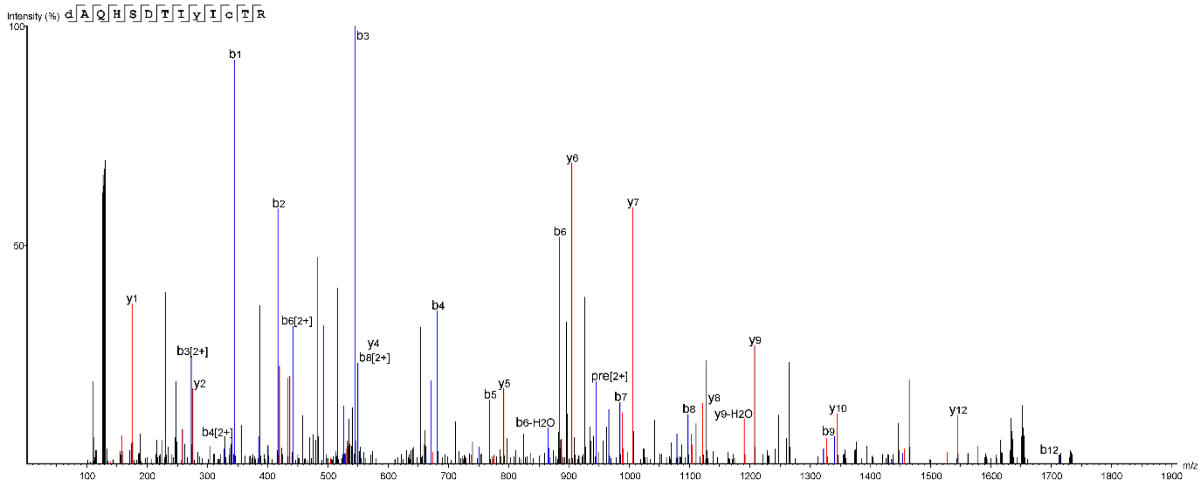


### AGC essential kinase 1 (AEK1)

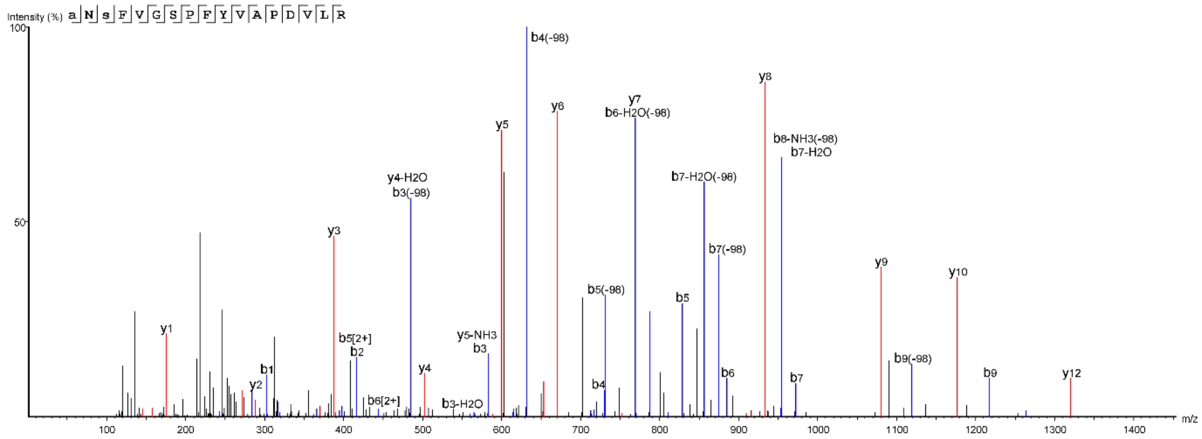

Mitogen-activated protein kinase kinase 5 (MKK5)

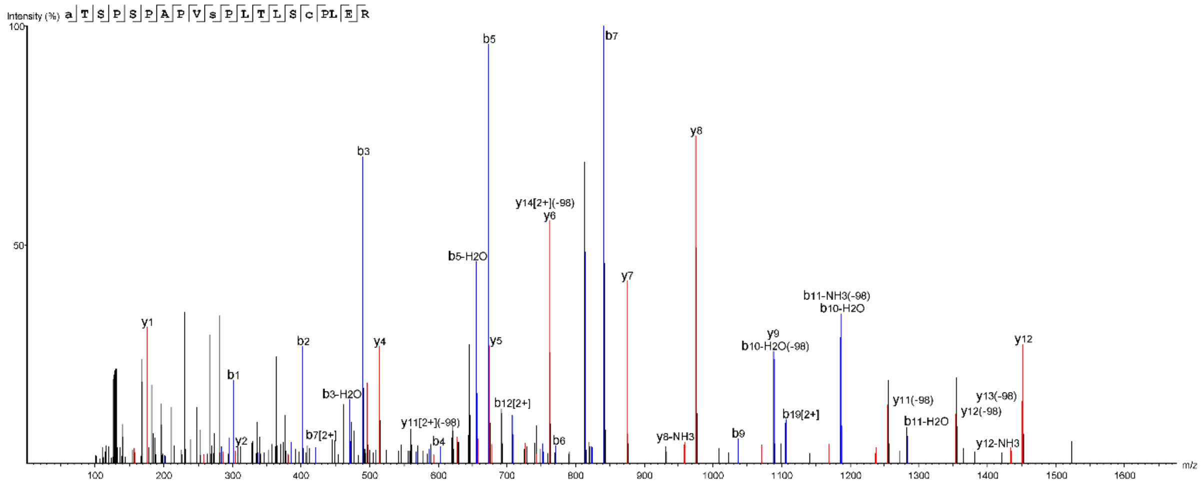


Mitogen-activated protein kinase kinase 7 (MKK7)

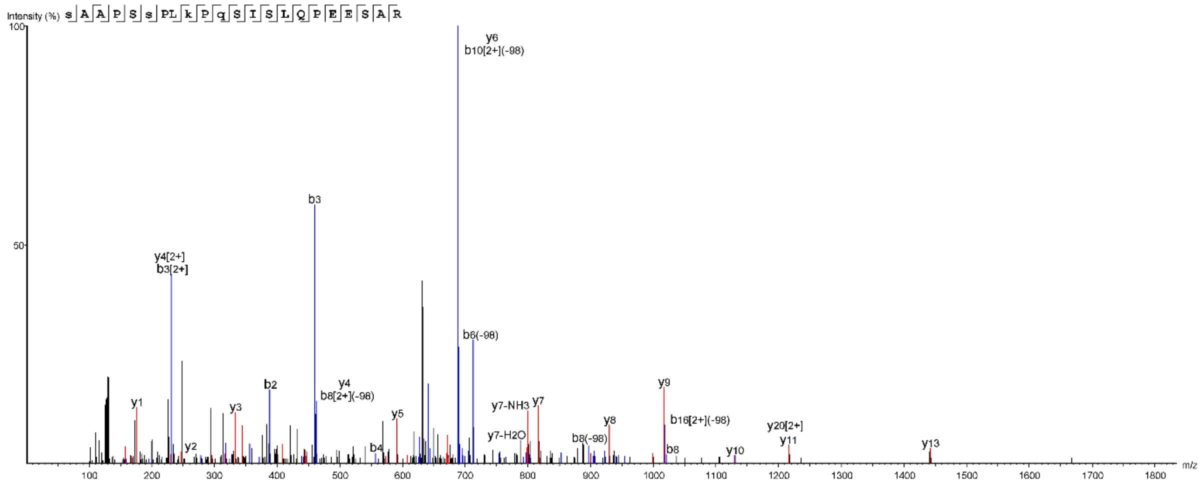


Glycogen synthase kinase 3 beta (GSK3β)

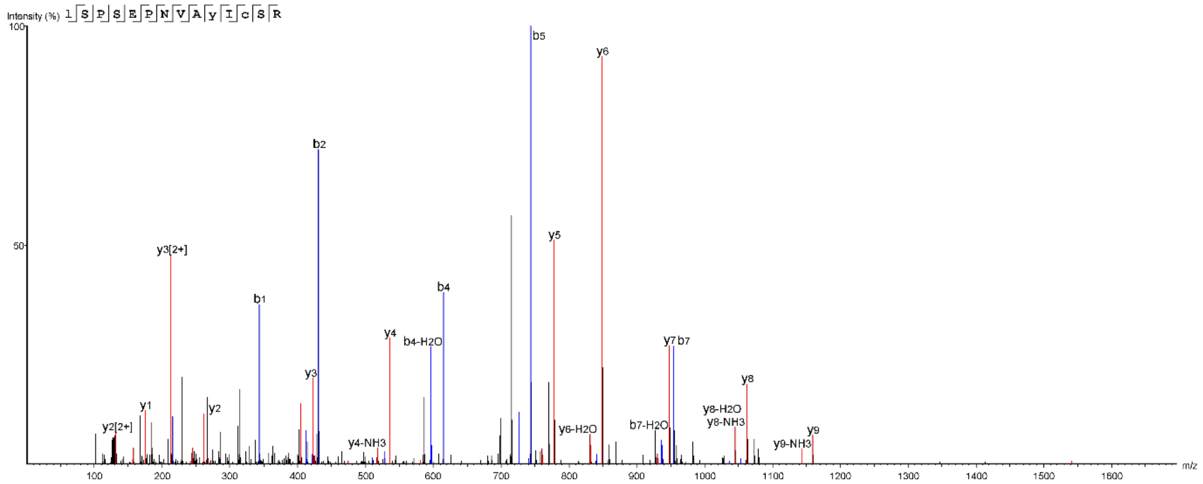

Glycogen synthase kinase 3 beta (GSK3β)

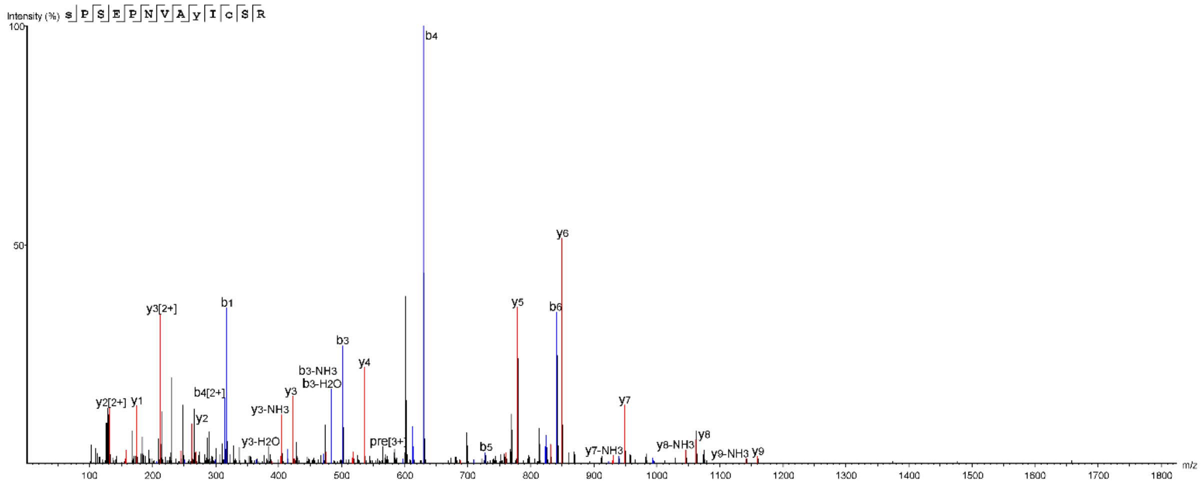

Glycogen synthase kinase 3 beta (GSK3β)

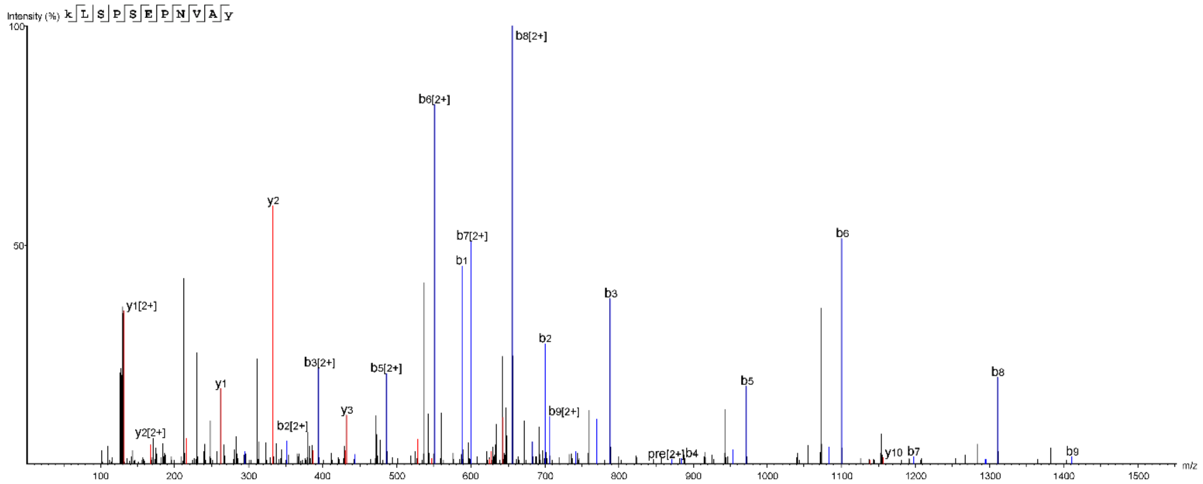

Protein kinase A (PKA)

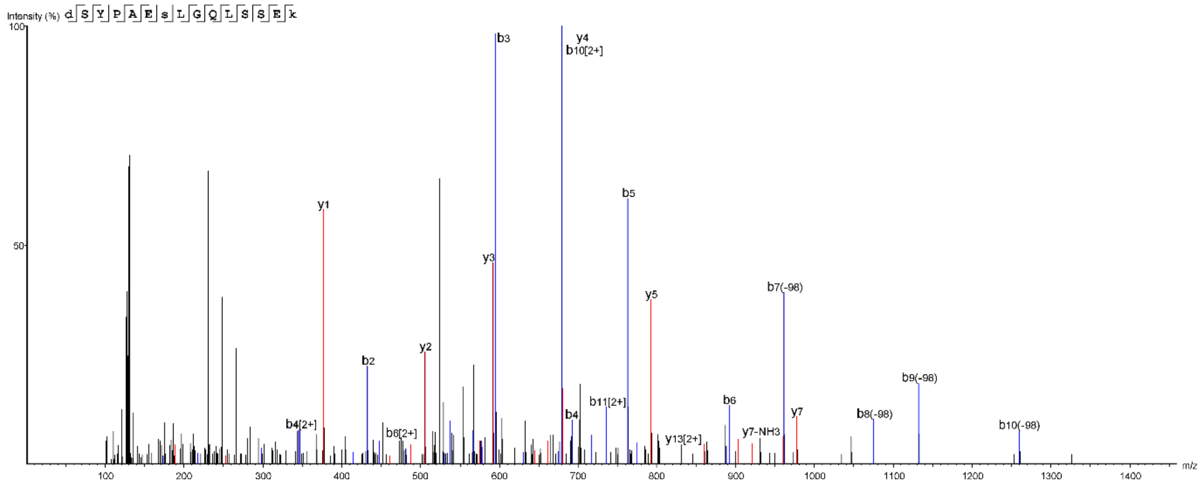

DEATH kinesin (LmxM.29.0350)

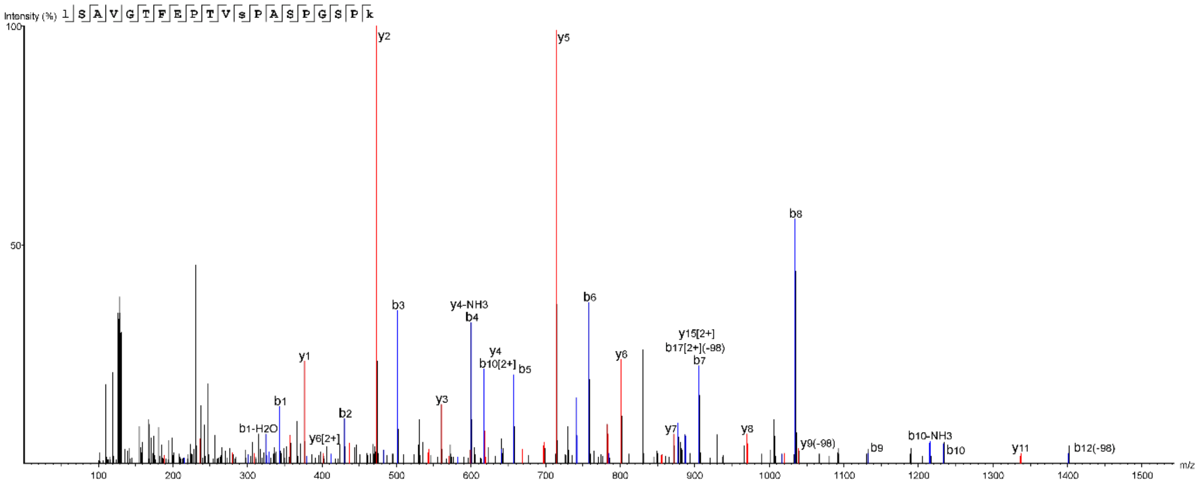


Kinesin D (LmxM.36.5150)

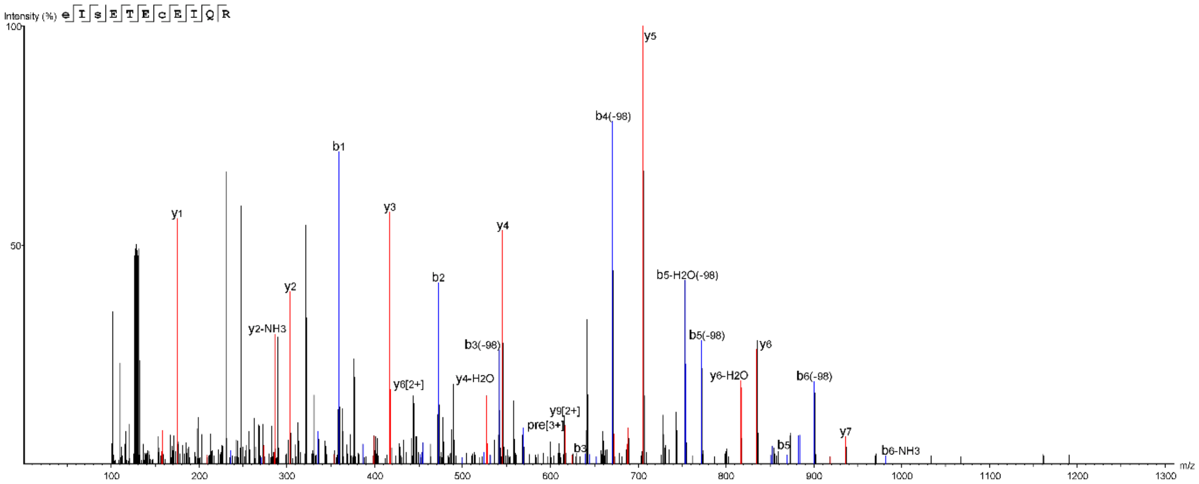


Kinesin (LmxM.11.0870)

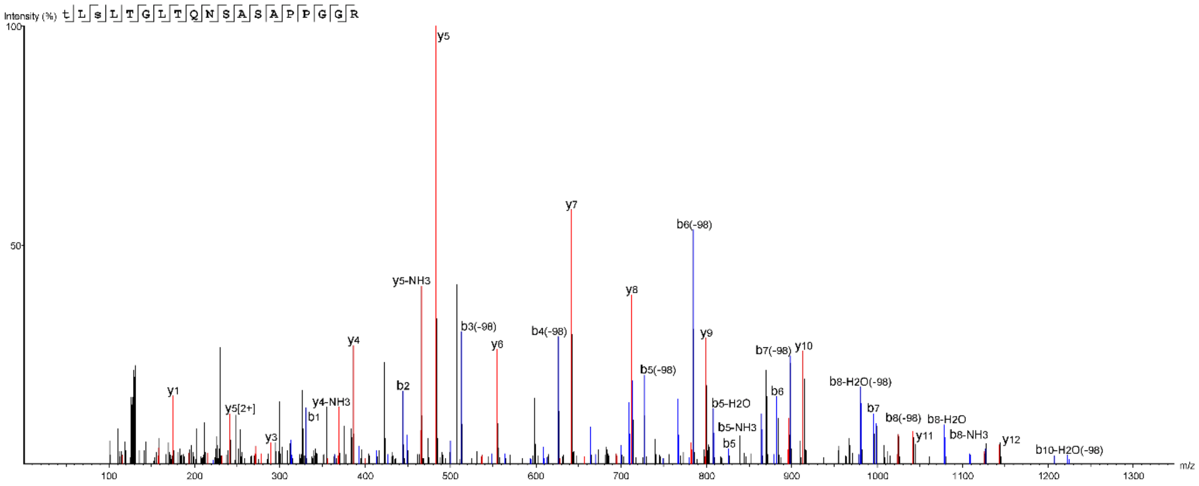

ADP/ATP mitochondrial carrier-like protein  (LmxM.14.0990)

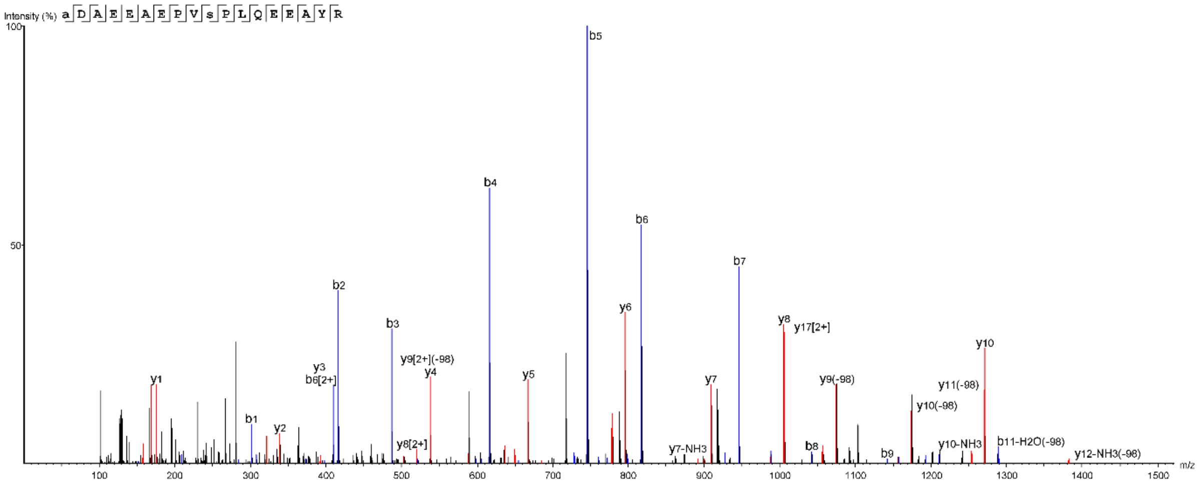


Isoprenylcysteine alpha-carbonyl methyltransferase (LmxM.15.0320)

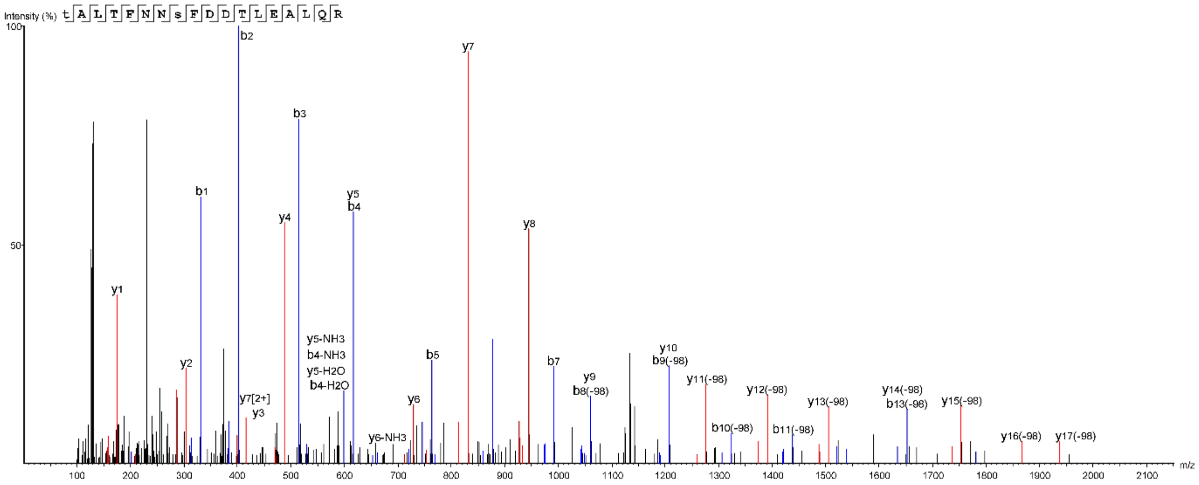


Carnosine N-methyltransferase (LmxM.33.1020)

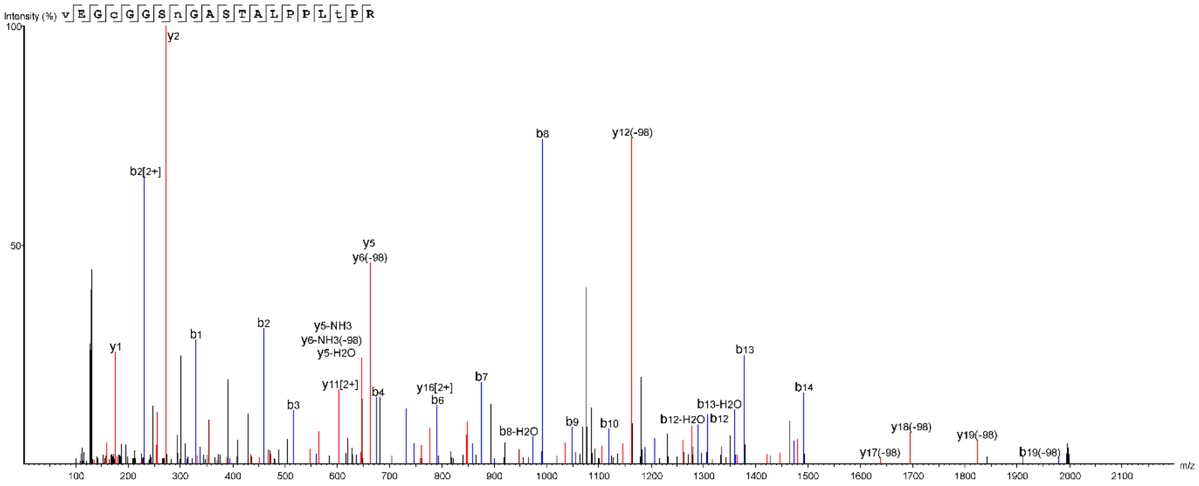

Distal docking complex protein 1 (LmxM.15.0540)

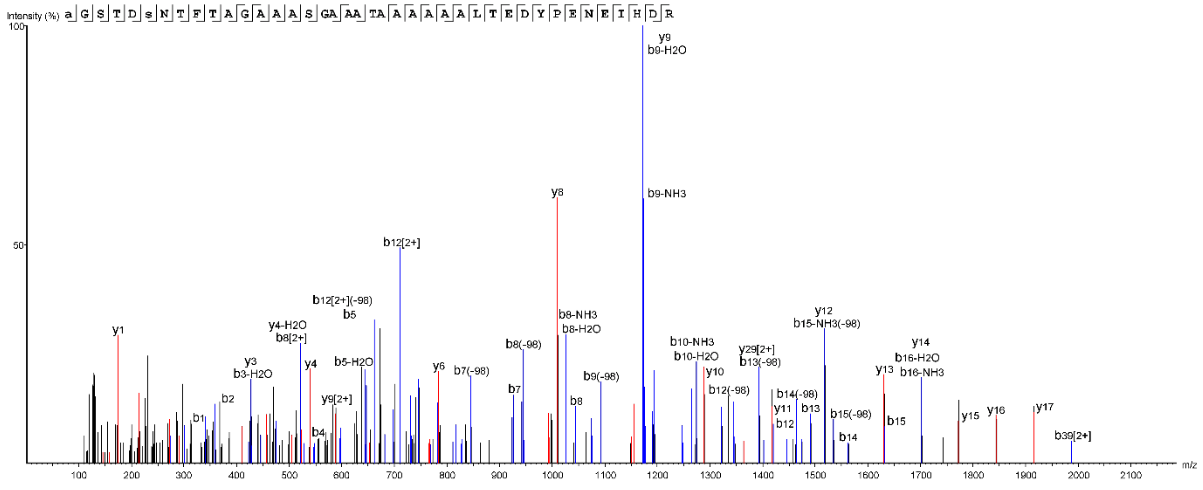

Cid1 family poly A polymerase (LmxM.19.1410)

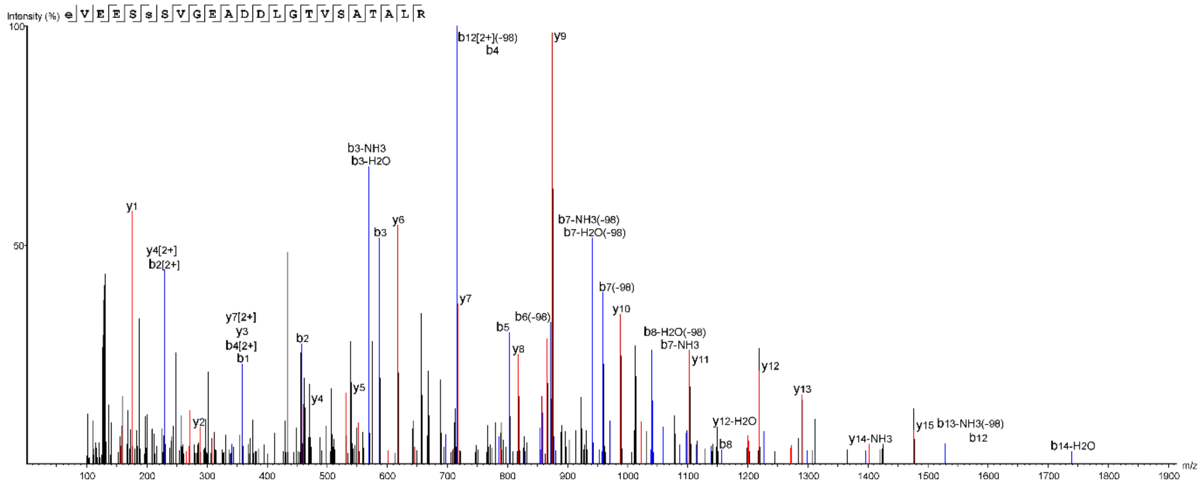

U3 small nucleolar ribonucleoprotein protein 10 (MPP10)

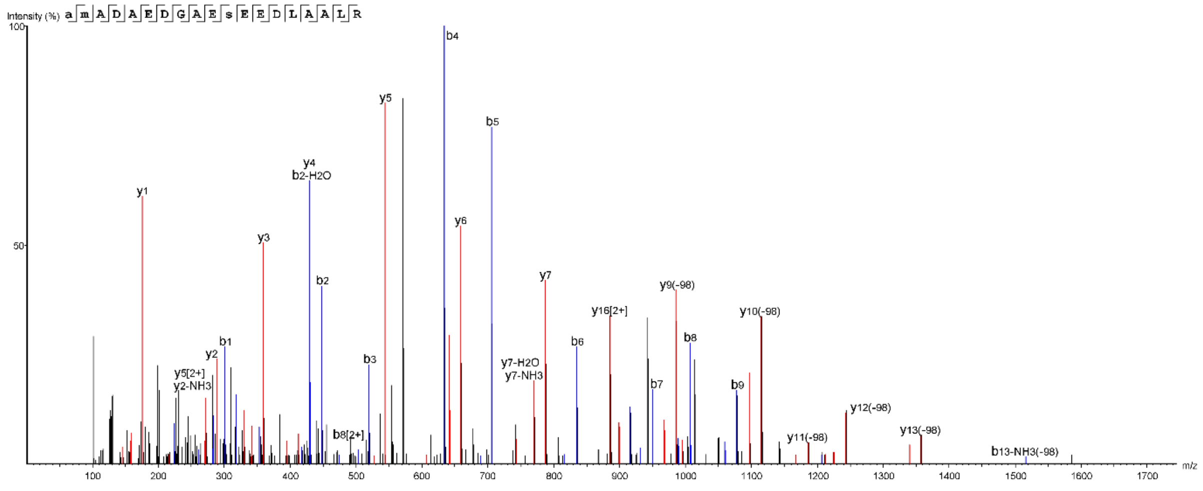

### DEAD/DEAH box helicase (LmxM.03.0690)

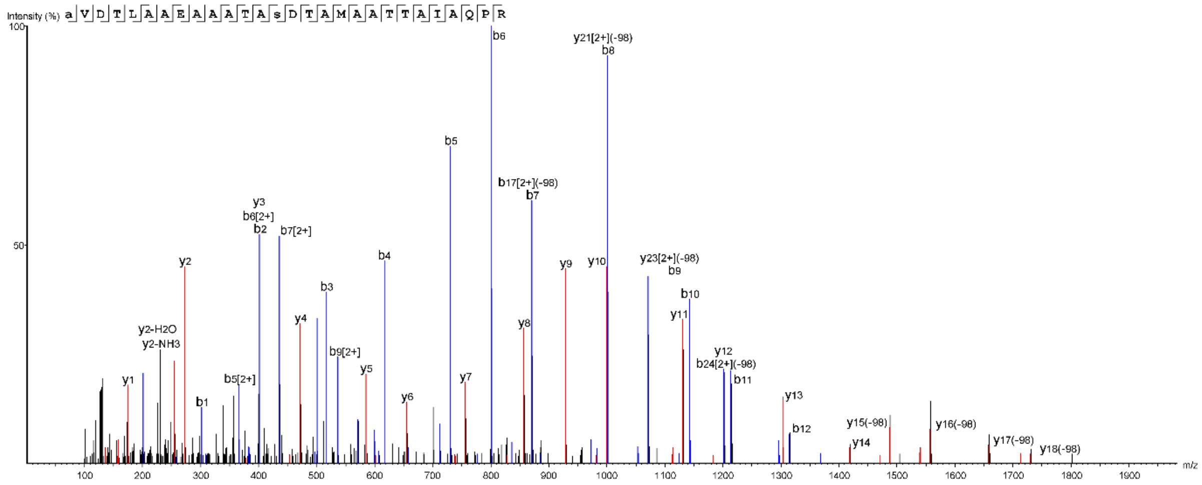


### ATP-dependent RNA helicase (HEL67)

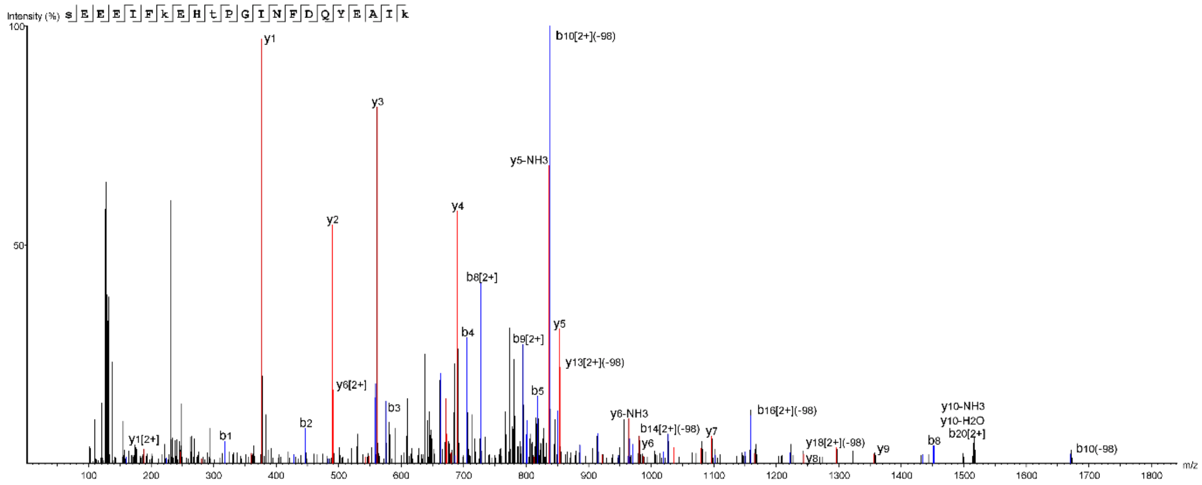


### ATP-dependent RNA helicase (LmxM.28.1310)

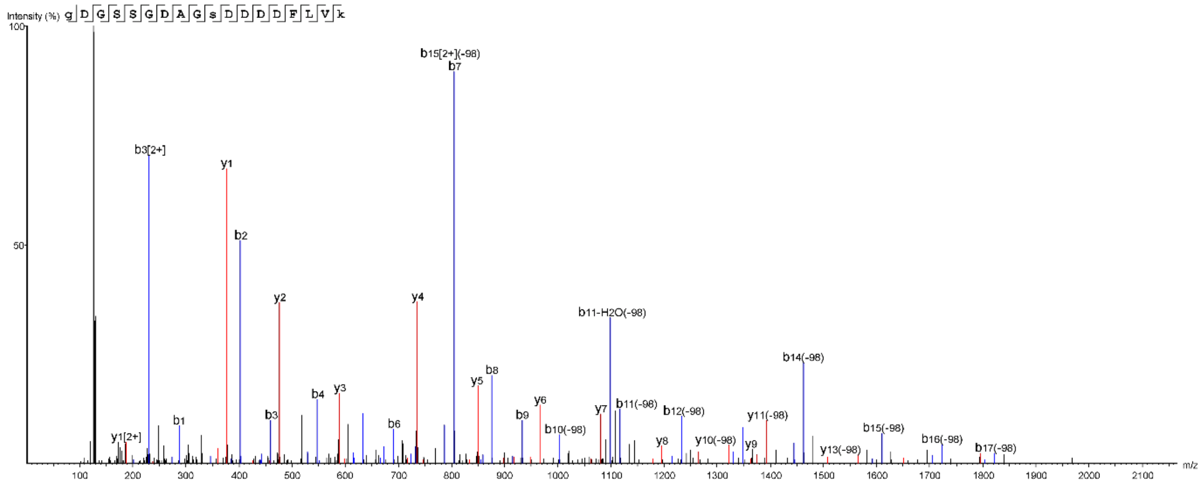

### KHARON (LmxM.36.5850)

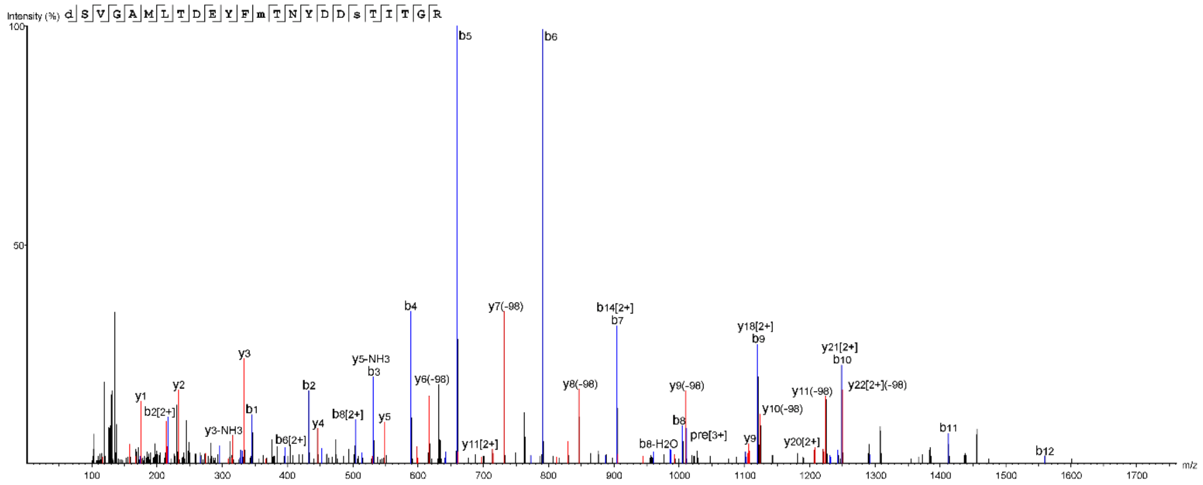


### Ubiquitin hydrolase (DUB3)

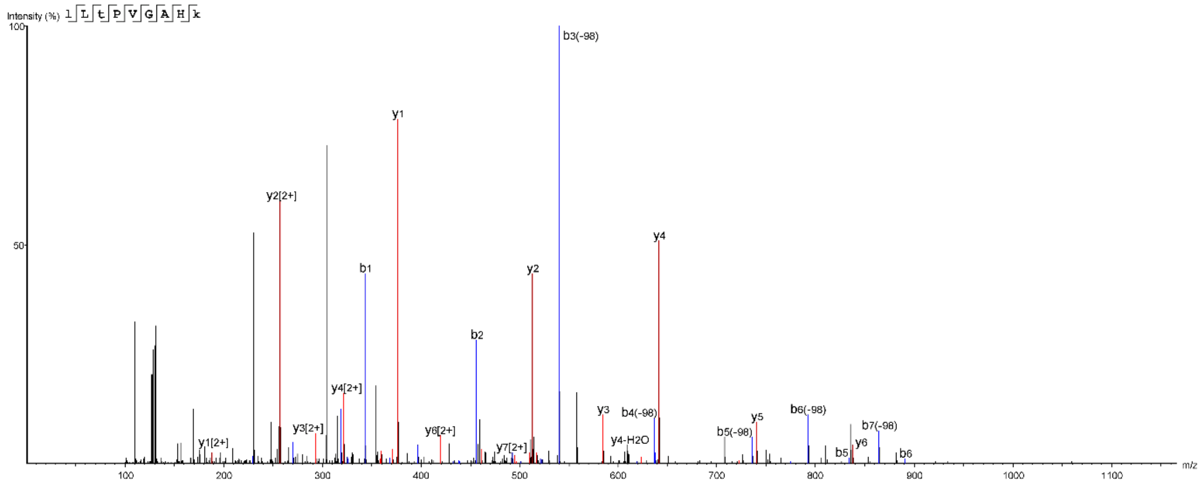


### Ubiquitin ligase (LmxM.09.0740)

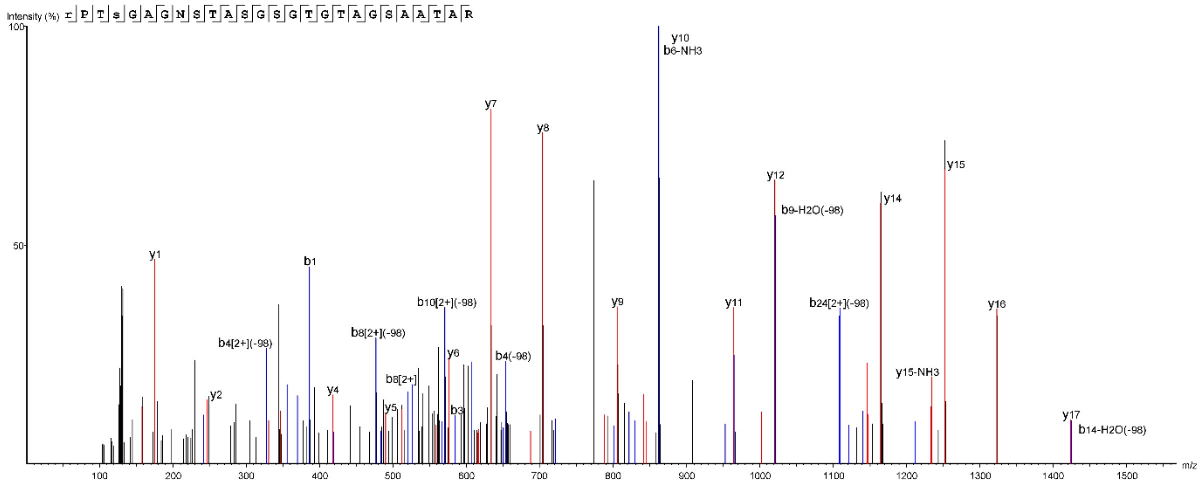

HECT E3 ligase (HECT3)

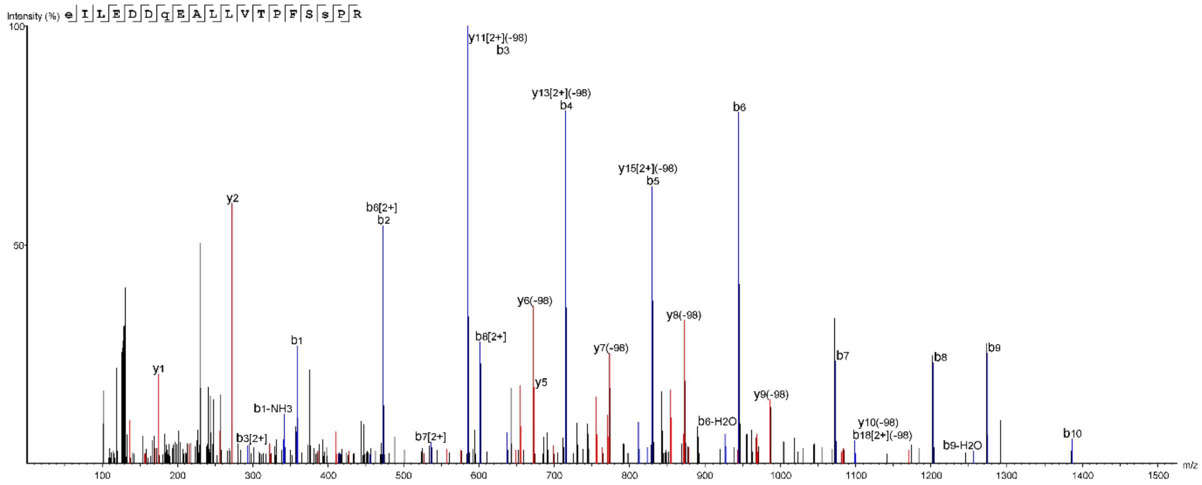


HECT E3 ligase (HECT7)

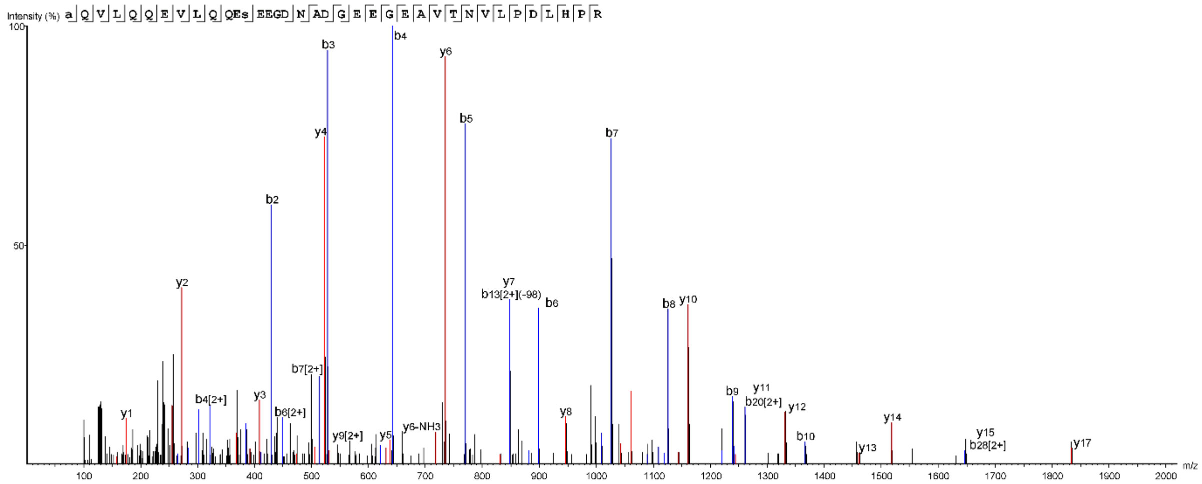


**FIG. S2** MS/MS spectra of phosphorylated tryptic peptides from selected *L. mexicana* proteins

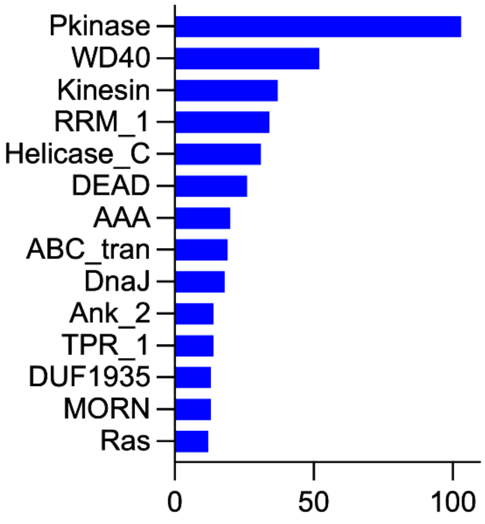


**FIG. S3** AXA vs LPP with Pfam annotation bar chart.

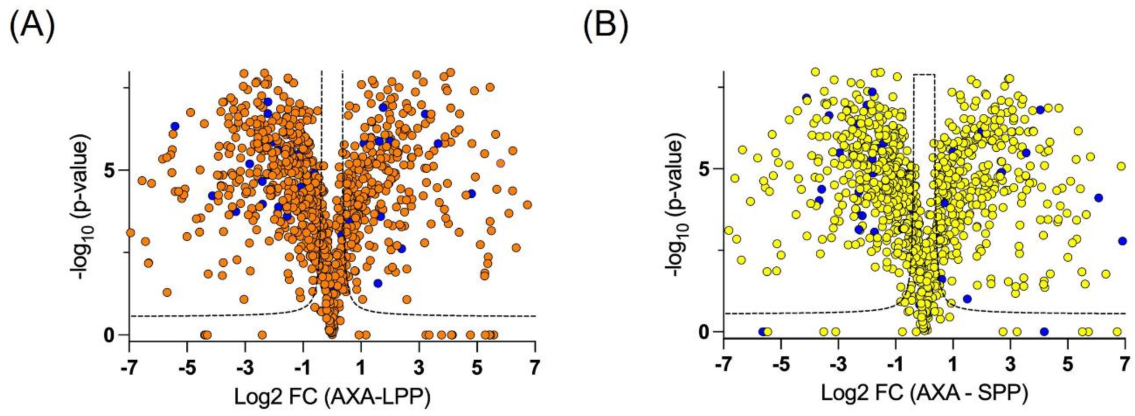


**FIG. S4** AXA vs LPP (A) and AXA vs SPP (B) volcano plots with kinases highlighted in blue.

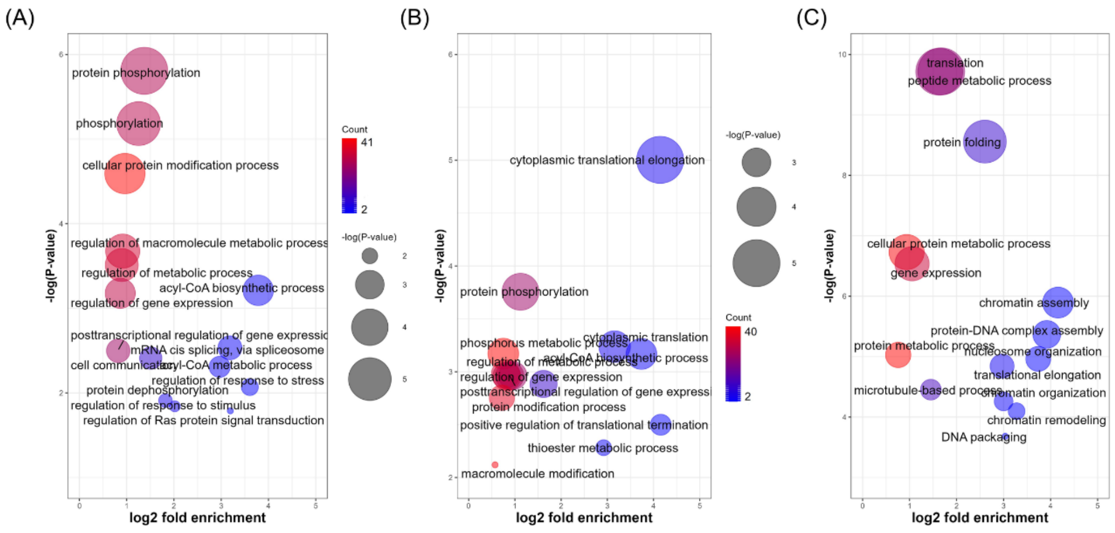


**FIG. S5** Biological Process (BP) gene ontology (GO) terms enriched in (A) LPP, (B) SPP and (C) AXA phosphoproteins.

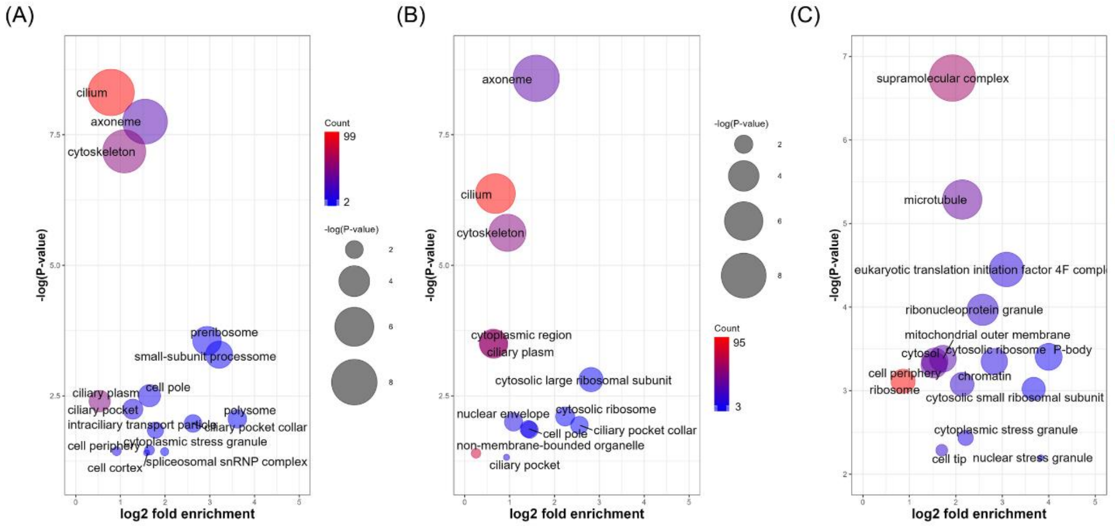


**FIG. S6** Cellular Component (CC) gene ontology (GO) terms enriched in (A) LPP, (B) SPP and (C) AXA phosphoproteins.

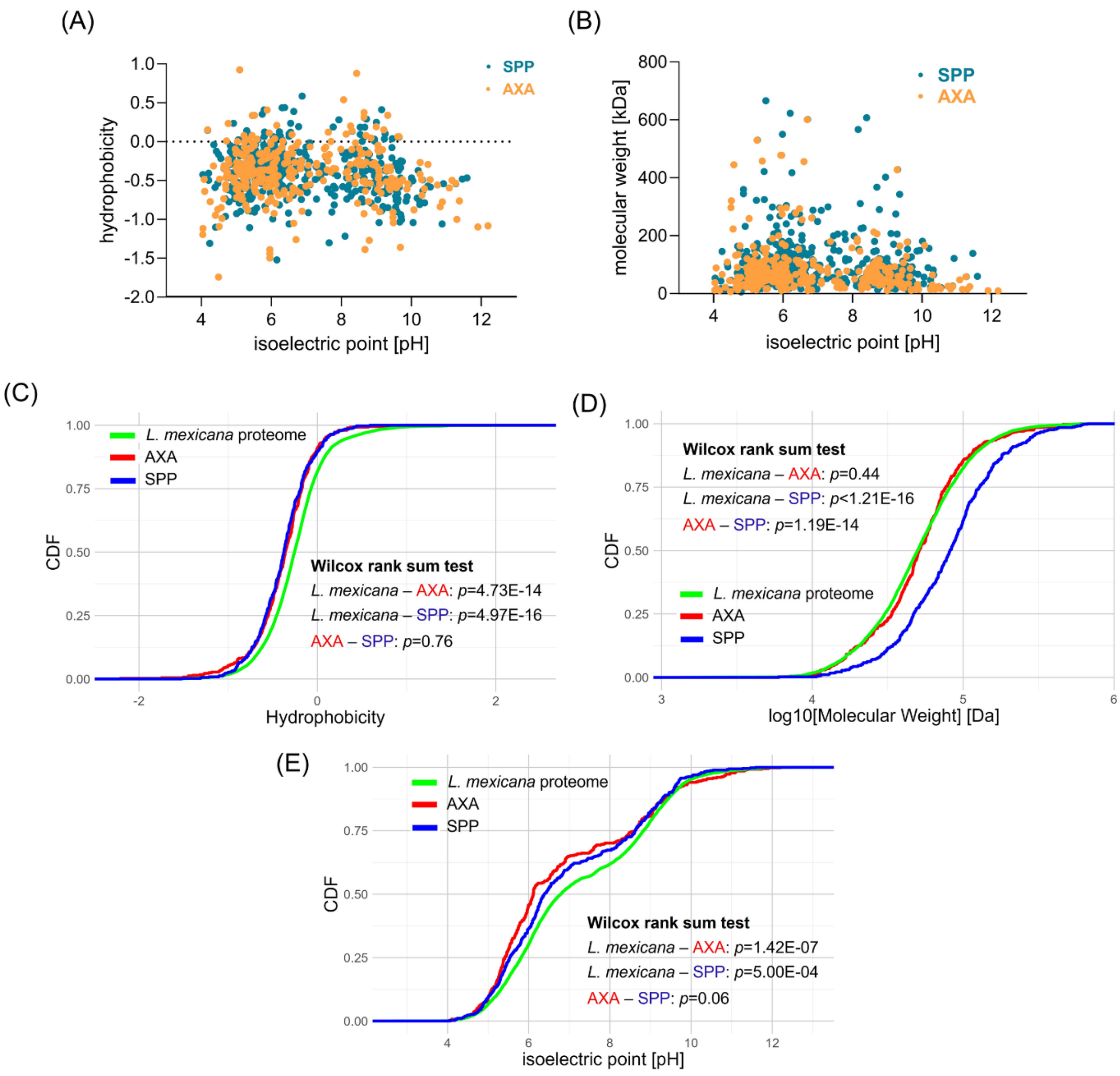


**FIG. S7** Physicochemical properties of *L. mexicana* phosphorylation substrates. (A) and (B) Scatter plots comparing hydrophobicity and isoelectric points and molecular weights and isoelectric points respectively of phosphorylation substrates in stationary phase promastigote (SPP) and axenic amastigote (AXA) life cycle stages. (C), (D) and (E) Cumulative distributions of hydrophobicity, molecular weights and isoelectric points respectively in the AXA and SPP phosphorylation substrates and the entire *L. mexicana* proteome. Wilcox rank sum test *p* values of the comparisons *L. mexicana* total proteome vs AXA phosphorylation substrates (*L. mexicana* – AXA) , *L. mexicana* total proteome vs SPP phosphorylation substrates (*L. mexicana* – SPP) and AXA vs SPP phosphorylation substrates (AXA – SPP) are shown.

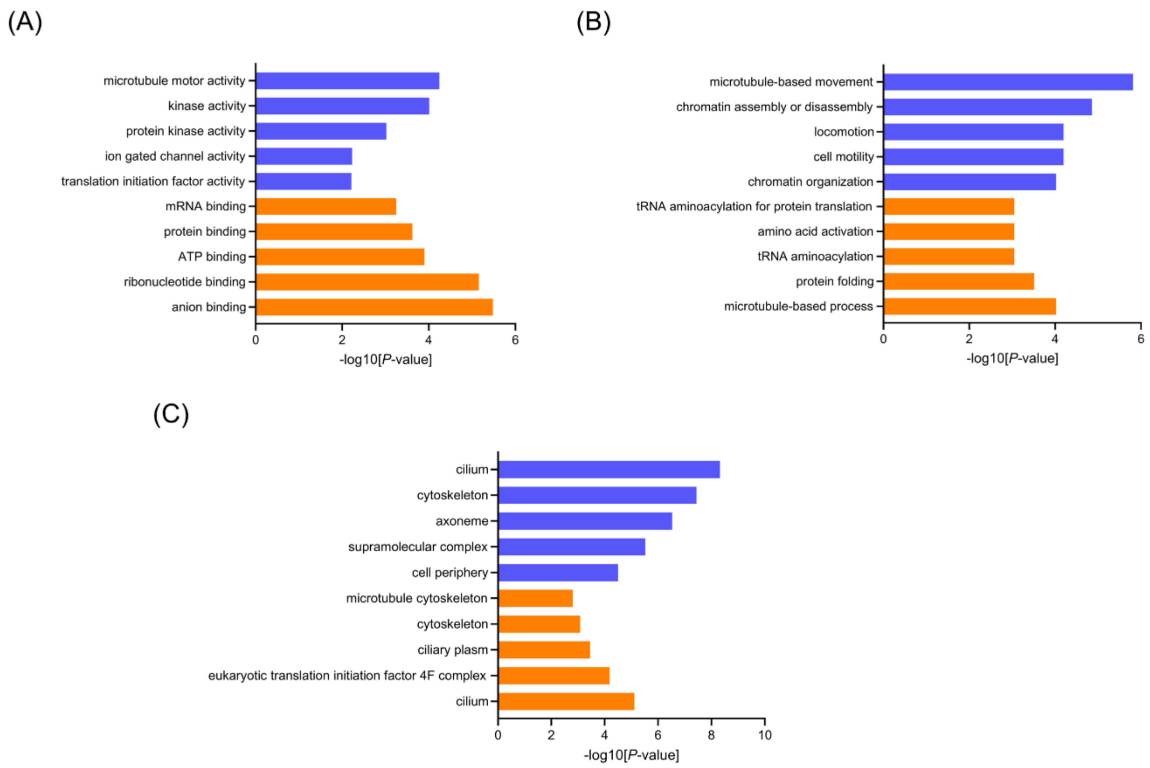


**FIG. S8** GO analysis of tanespimycin-affected phosphoproteins in *L. mexicana* log phase promastigotes (LPP). (A), (B) and (C) are Molecular Function (MF), Biological Process (BP) and Cellular Component (CC) GO terms respectively; blue: negatively affected phosphoproteins, red: positively affected phosphoproteins.

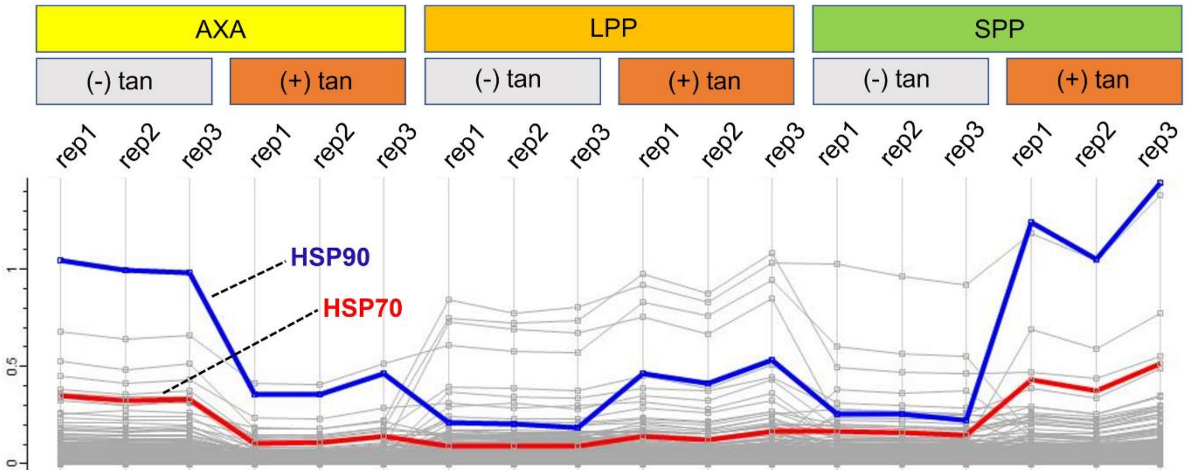


**FIG. S9** Profile plot of the *L. mexicana* phosphoproteins with (+) and without (-) tanespimycin (tan) treatment in axenic amastigote (AXA), log phase promastigote (LPP) and stationary phase promastigote (SPP) life cycle stages. HSP90 (blue) and HSP70 (red) proteins are highlighted.

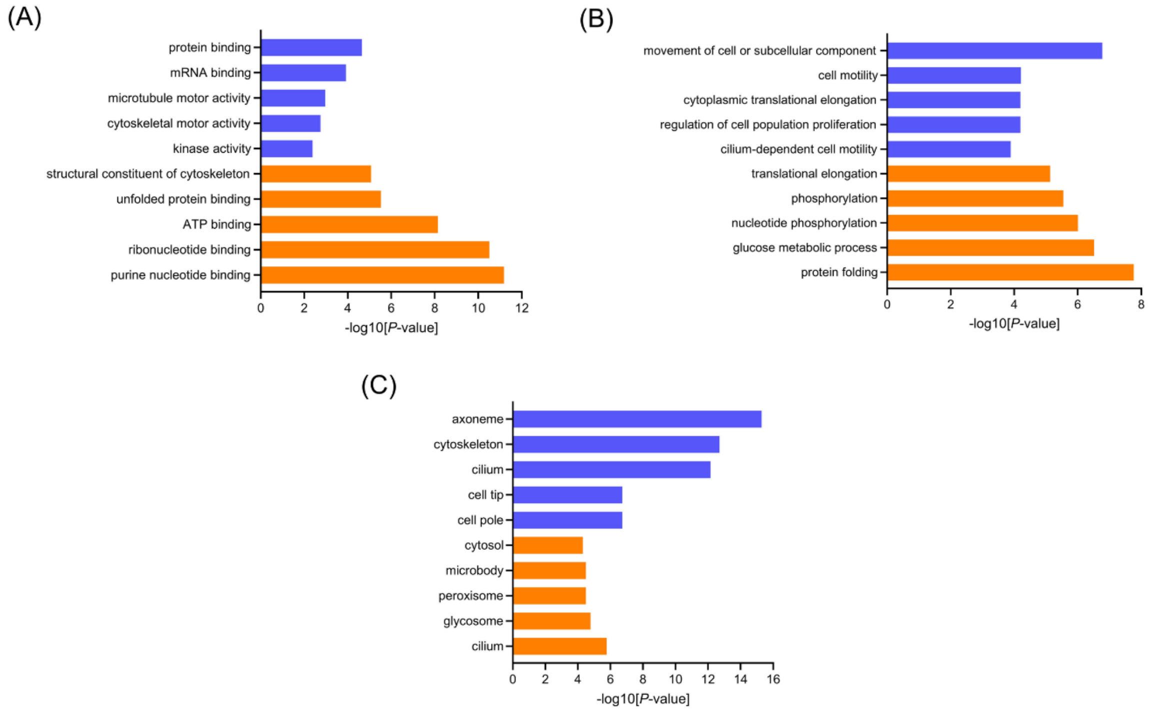


**FIG. S10** GO analysis of tanespimycin-affected phosphoproteins in *L. mexicana* stationary phase promastigotes (SPP). (A), (B) and (C) are Molecular Function (MF), Biological Process (BP) and Cellular Component (CC) GO terms respectively; blue: negatively affected phosphoproteins, red: positively affected phosphoproteins.

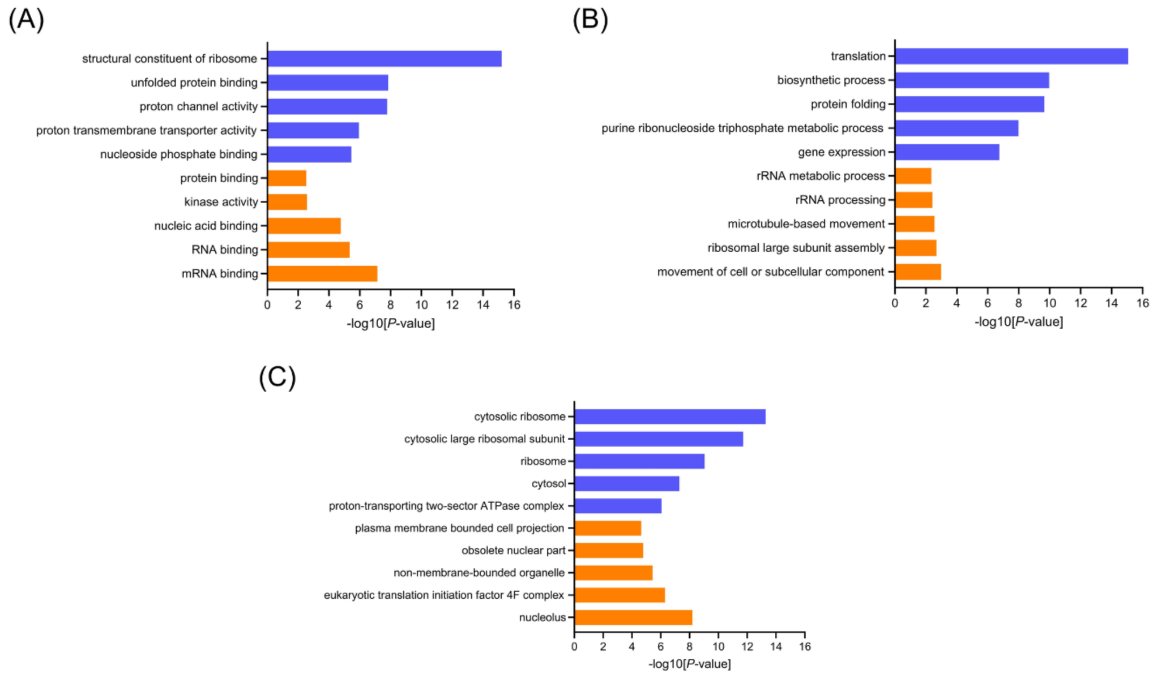

**FIG. S11** GO analysis of tanespimycin-affected phosphoproteins in *L. mexicana* axenic amastigotes (AXA). (A), (B) and (C) are Molecular Function (MF), Biological Process (BP) and Cellular Component (CC) GO terms respectively; blue: negatively affected phosphoproteins, red: positively affected phosphoproteins.

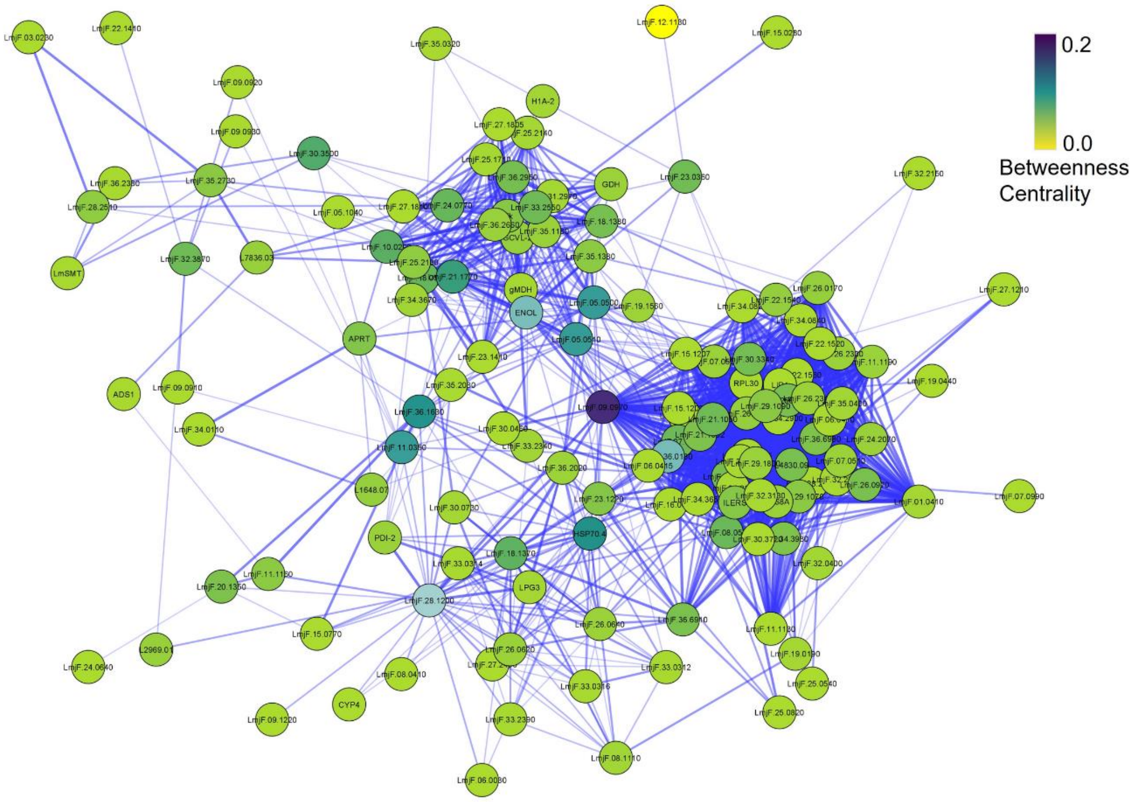


**FIG. S12** Protein-protein interaction network of RNA-binding phosphoproteins downregulated during HSP90 inhibition in *L. mexicana* axenic amastigotes (AXAs) constructed using publicly available STRING database of *L. major* Friedlin strain. The nodes are coloured according to their betweenness centrality in the network.

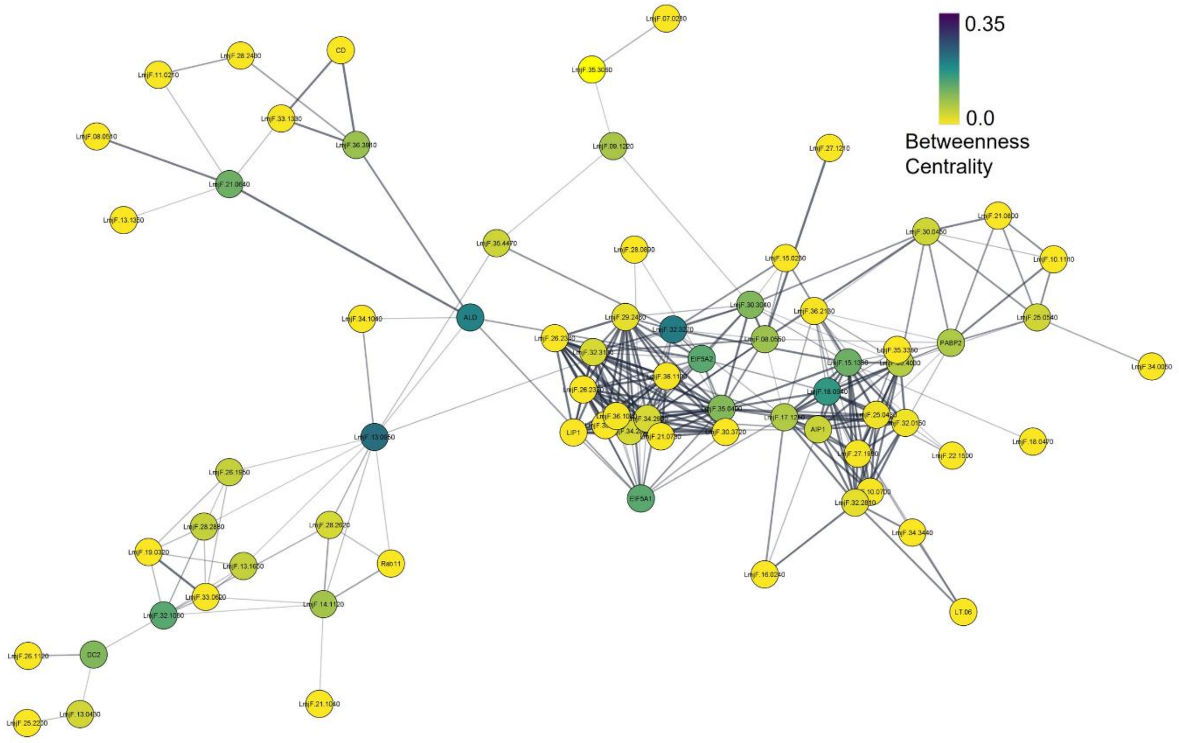


**FIG. S13** Protein-protein interaction network of RNA-binding phosphoproteins downregulated during HSP90 inhibition in *L. mexicana* log phase promastigotes (LPPs) constructed using publicly available STRING database of *L. major* Friedlin strain. The nodes are coloured according to their betweenness centrality in the network.

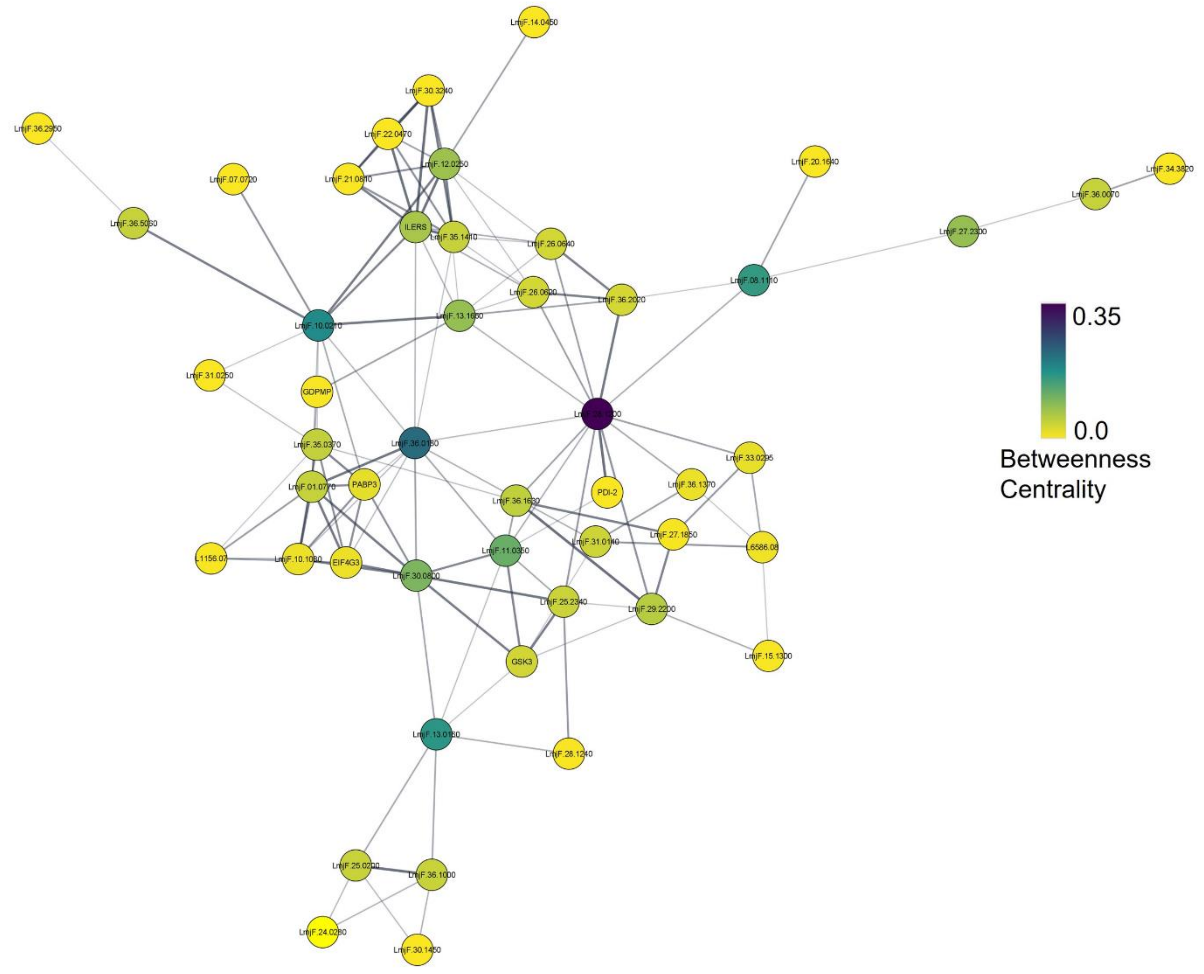


**FIG. S14** Protein-protein interaction network of RNA-binding phosphoproteins upregulated during HSP90 inhibition in *L. mexicana* log phase promastigotes (LPPs) constructed using publicly available STRING database of *L. major* Friedlin strain. The nodes are coloured according to their betweenness centrality in the network.
